# Electron transport chain complex I and mitochondrial fusion regulate ROS for differentiation in *Drosophila* neural stem cells

**DOI:** 10.1101/2025.07.22.666060

**Authors:** Rahul Kumar Verma, Atharva Bhingare, Dnyanesh Dubal, Richa Rikhy

**Affiliations:** Biology, Indian Institute of Science Education and Research, Homi Bhabha Road, Pashan, Pune, 411008, India; Laboratoire de Biologie et Modélisation de la Cellule, Univ Lyon, ENS de Lyon, UCB Lyon 1, CNRS, Lyon, France; BESE, KAUST, Thuwal, Kingdom of Saudi Arabia

**Keywords:** neural stem cells, *Drosophila*, electron transport chain, complex 1, mitochondria, ROS, cyclin E, Drp1

## Abstract

Mitochondrial activity and dynamics play crucial roles in regulating neuronal differentiation. Neural stem cells (NSCs) or neuroblasts in *Drosophila* require the regulation of mitochondrial fusion, along with the activity of the electron transport chain (ETC), to meet the metabolic demands of differentiation. However, the mechanisms by which mitochondrial fusion and activity together regulate NSC differentiation are not well understood. We investigated the relationship between mitochondrial fusion, and ETC complex I activity during *Drosophila* neuroblast differentiation. We found that depletion of complex I subunits did not affect the number of type II neuroblasts but reduced their proliferation, thereby decreasing the numbers of mature intermediate precursor cells (mINPs), ganglion mother cells (GMCs), and neurons in each lineage. Complex I depletion decreased the mitochondrial membrane potential and cristae numbers, and increased mitochondrial fragmentation and ROS. Increased ROS resulting from the depletion of antioxidant enzymes also led to a decrease in mINPs, GMCs, and neurons in each type II neuroblast lineage. Both complex I and antioxidant protein depletion led to delayed G1/S transition and decreased nuclear cyclin E levels. Interestingly, the defects in proliferation, differentiation, and ROS in complex I and anti-oxidant protein depleted neuroblasts could be restored by fused mitochondrial morphology obtained through additional depletion of the fission protein Drp1. Further, overexpression of anti-oxidant proteins could alleviate the ROS and rescue the differentiation defect in complex I depleted type II neuroblasts. Together, this study reveals a role for complex I and mitochondrial fusion in restricting ROS for differentiation in *Drosophila* neuroblasts.

**Significance statement:** The electron transport chain complex I regulates differentiation in the *Drosophila* type II neuroblast lineage. Complex I and anti-oxidant scavenger protein depletion lead to mitochondrial fragmentation and an increase in ROS, thereby affecting the G1/S phase transition and decreasing the formation of lineage cells. Mitochondrial fusion induced by depletion of mitochondrial fission protein Drp1 in complex I and anti-oxidant protein depleted type II neuroblasts leads to suppression of the differentiation defects. This study confirms a crucial role of mitochondrial fusion in restricting ROS for appropriate division and differentiation in the type II neuroblast lineage in *Drosophila*.

## Introduction

Mitochondrial dynamics and activity regulate stem cell differentiation. In mammals, pluripotent stem cells predominantly rely on glycolysis for energy production. They contain fragmented or globular mitochondria, with poorly organised cristae and perinuclear localisation. These fragmented mitochondria have low electron transport chain (ETC) activity, decreased mitochondrial membrane potential (MMP) and ATP production. During differentiation, mitochondria transition to oxidative phosphorylation (OXPHOS), facilitated by fused or tubular mitochondrial architecture with well-developed cristae. Fused mitochondria have high OXPHOS, MMP, and ATP production (1–4). An equilibrium between the fusion and fission events determines the mitochondrial architecture and consequently, their activity. The family of large Dynamin GTPases orchestrates these events. Mitofusin (Mfn) in mammals, or Mitochondrial assembly regulatory factor (Marf) in *Drosophila,* drives outer mitochondrial membrane fusion, followed by inner mitochondrial membrane fusion through Optic atrophy 1 (Opa1). Dynamin-related protein 1 (Drp1) is essential for mitochondrial fragmentation.

Mitochondrial dynamics and activity regulate proliferation and differentiation in *Drosophila* neural stem cells, also known as neuroblasts. In larval stages, proliferating neuroblasts rely on glycolysis; however, during the larval-to-pupal transition, they exit the cell cycle and undergo terminal differentiation (5).

Mitochondrial fusion driven by Opa1 is essential for differentiation in the central brain neuroblasts (6). An increase in Notch signaling leads to excessive proliferation of neuroblasts, which also requires mitochondrial activity and fusion (7, 8). Although mitochondrial dynamics and activity are crucial for neuroblast proliferation and differentiation, the mechanisms by which they interact to produce differentiation outcomes remain largely unexplored.

In this study, we investigate the interaction between mitochondrial fusion and oxidative phosphorylation driven by complex I during the proliferation and differentiation of neuroblasts in the central brain of *Drosophila*. The central brain in the third instar larva contains type I and type II neuroblasts. The type I neuroblasts are approximately 90 in number, and the type II neuroblasts are 8 in number. They express transcription factors Deadpan (Dpn) and Asense (Ase). Type I neuroblasts divide to give rise to ganglion mother cells (GMCs). The type II neuroblast lineages contain transit amplifying cells, similar to those found in mammalian systems. Type II neuroblasts express an Ets transcription factor, *pointed P1* (*pnt*) (9). They undergo division to produce immature intermediate neural progenitors (INPs) (10). Immature INPs undergo transcriptional changes to form mature INPs (mINPs), which express Dpn and Ase. mINPs divide asymmetrically 5–6 times, to form GMCs. GMCs in both type I and II lineages express Prospero (Pros) and Ase. GMCs eventually divide symmetrically to produce neurons or glia (Fig. 1*A*). High nuclear Pros and Elav are expressed in neurons (11, 12).

**Fig. 1.**
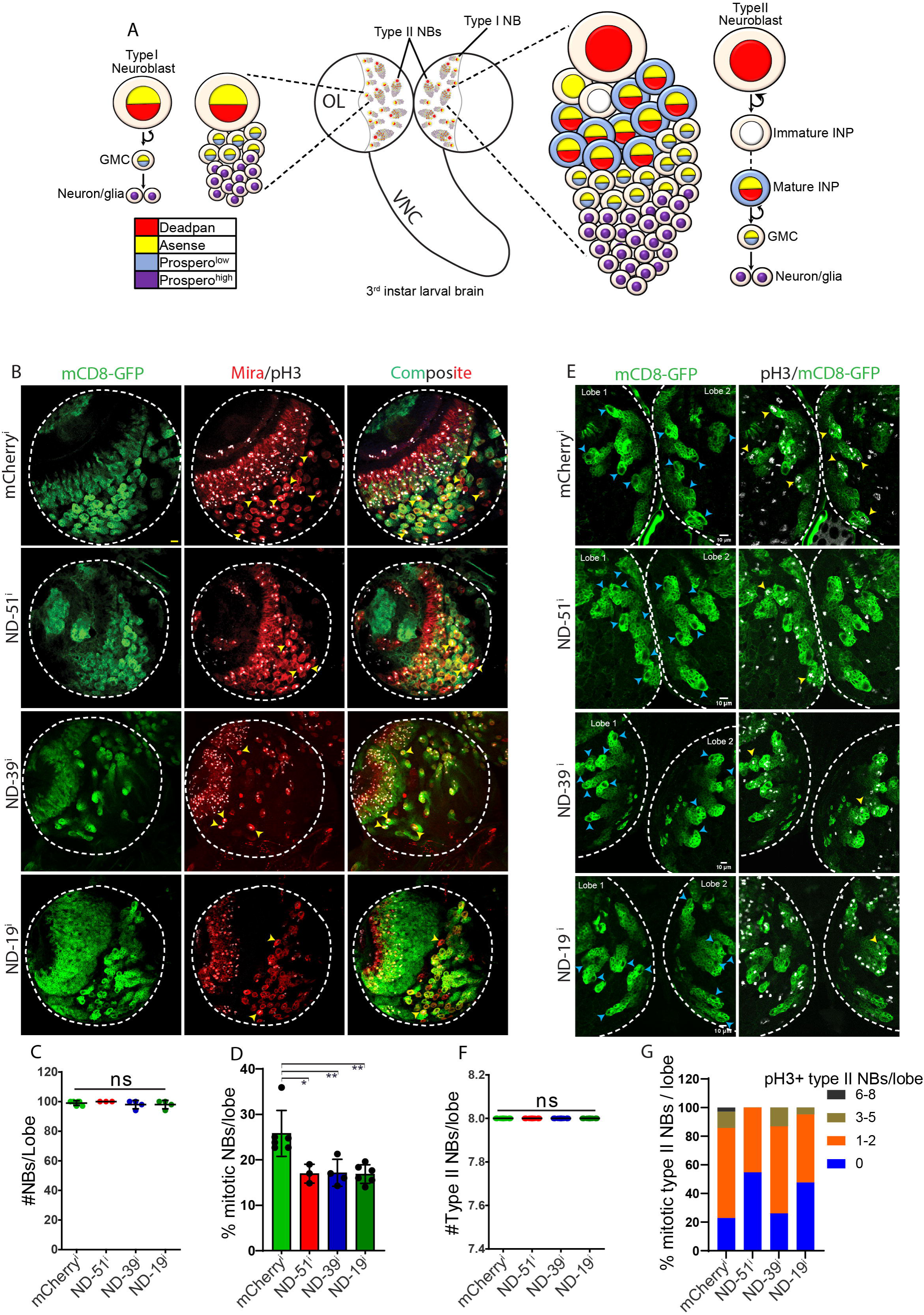
Deletion of complex I reduces the proliferation of neural stem cells in the third instar larval brain. **(A)** The *Drosophila* third instar larval brain contains type I and type II neuroblasts, which divide asymmetrically. Type I neuroblasts divide symmetrically to self-renew and produce one GMC, which divides once to give two neurons or glia. Type II neuroblasts self-renew and generate one immature INP. After a maturation phase, INPs divide 4-5 times asymmetrically to self renew and generate GMCs, which then divide symmetrically to form neurons or glia. Distinct cellular markers label each progenitor population: Red-Dpn, Yellow-Ase, Blue-Low cytoplasmic Pros, Purple-High nuclear Pros. **(B)** Representative images for mitotically dividing neural stem cells (NSCs) (Yellow arrowheads) in the *Drosophila* third instar larval brain lobe of indicated genotypes. *wor*-Gal4 driven UAS-mCD8-GFP marks the boundary of stem cells and its lineage (green). Miranda labels the stem cell cortex (red) and mitotically dividing cells are labelled by pH3 (grey). Scale bar-10µm. **(C)** Graph showing the quantification of percent mitotically dividing NSCs present in each brain lobe of third instar larvae, expressing *wor*-Gal4 driven mCherry^i^ (n=607 NSCs, 6 brains), ND-51^i^ (301, 3), ND-391^i^ (473, 5), and ND-19^i^ (605, 6). **(D)** Representative images of mitotically dividing mINPs in type II neuroblasts (Yellow arrowheads) in the *Drosophila* third instar larval brain of indicated genotypes. *pnt*-Gal4 driven mCD8-GFP marks the boundary of stem cells (blue arrowheads) and its lineage (green), and mitotically dividing cells are labelled by pH3 (grey). Scale bar-10µm. **(E)** Quantification of mitotically dividing type II neuroblasts expressing *pnt*-Gal4 driven mCherry^i^ (n=144 neuroblasts, 9 brains), ND-51^i^ (176, 11), ND-39^i^ (192, 12), and ND-19^i^ (176, 11).

ETC complex I, known as NADH dehydrogenase or NADH:ubiquinone oxidoreductase is a multi-subunit enzyme with a 1000 kDa molecular weight that catalyzes the transfer of electrons from NADH to ubiquinone. Complex I can generate ROS during electron transfer in the ETC. It is an L-shaped structure, consisting of one hydrophobic arm embedded in the inner mitochondrial membrane and one hydrophilic arm present at the periphery, facing the matrix (13–15). The *Drosophila* complex I is composed of 43 subunits, a catalytic core of 14 evolutionarily conserved subunits alongside 29 accessory subunits (13, 14). Although these accessory subunits are not directly involved in the catalytic process, their perturbation can disrupt the assembly of the entire complex, leading to elevated ROS generation (16, 17). Electrons from complex I are transferred to complex III and then to IV, where molecular oxygen is reduced to water. This electron flow is coupled to the pumping of protons from the mitochondrial matrix to the intermembrane space by complex I, II, and IV, establishing an electrochemical gradient across the inner mitochondrial membrane, which manifests as the MMP. Complex V (ATP Synthase) utilises the MMP to synthesise ATP from ADP.

In this study, we found that depletion of complex I subunits led to decreased proliferation in the central brain neuroblasts. There was a decrease in differentiation in the type II neuroblast lineage, characterized by reduction in the numbers of mINPs, GMCs, and neurons. Notably, the downregulation of complex I led to fragmented mitochondria, decreased cristae numbers, a low MMP, and increased ROS generation. These elevated ROS levels led to a decrease in differentiation by disrupting the transition from the G1 to the S phase in the cell cycle. Inducing mitochondrial fusion or increasing ROS scavengers in complex I depleted type II neuroblasts reversed the ROS levels and restored the differentiation defects in the type II neuroblast lineage.

## Results

### ETC complex I depletion leads to a decrease in the proliferation and differentiation of neuroblasts at the larval stage

To evaluate the function of ETC complex I, we depleted individual subunits in type I and II neuroblasts by combining *worniu*-Gal4 (*wor-*Gal4), mCD8-GFP and *pointed P1*-Gal4 (*pnt*-Gal4), mCD8-GFP with shRNA lines against them (Fig. S1*A*). mCD8-GFP marked the plasma membrane in the cells expressing the Gal4 and the shRNA. RNAi depletion was carried out against the complex I subunits, ND-49 (mammalian NDUFS2) and ND-51 (NDUFV1), belonging to the core hydrophilic matrix arm and ND-19 (NDUFA8), ND-39 (NDUFA9), ND-42 (NDUFA10) and ND-PDSW (NDUFB10) subunits belonging to the accessory hydrophobic membrane arm. When expressed with *wor*-Gal4 at 29°C, we observed pupal lethality on the depletion of ND-PDSW, ND-49, and ND-42, with the few flies that emerged from pupae being sluggish in behaviour. Depletion of ND-51, ND-39, and ND-19 gave a more severe phenotype of complete pupal lethality (Fig. S1*A*). Depletion of complex I subunits specifically in type II neuroblasts using *pnt*-Gal4 did not affect the development of F1 flies. Flies emerged normally out of pupae but were less active and sluggish. They died within 15 days after eclosion (Fig. S1*A*). The shRNA lines against ND-51 (ND-51^i^), ND-39 (ND-39^i^), and ND-19 (ND-19^i^) were selected for further analysis of the role of complex I in neuroblast proliferation and differentiation.

We assessed the impact of ND-51, ND-39 and ND-19 depletion on the numbers of central brain neuroblasts and their proliferation by crossing the corresponding shRNA lines with *wor*-Gal4 and immunostaining for neuroblasts with Miranda and phospho-Histone 3 (pH3), which marks dividing cells. There was no significant difference in the numbers of neuroblasts between the controls and brains depleted of ND-51, ND-39, and ND-19 (Fig. 1*B* and *C*). However, there was a significant reduction in the proportion of mitotically active neuroblasts within each brain lobe (Fig. 1*B* and *D*). Knockdown of ND-51, ND-39, and ND-19 in type II neuroblasts using the *pnt*-Gal4 driver also yielded a decreased number of mitotically dividing neuroblasts as compared to controls, but the number of neuroblasts remained unchanged (Fig. 1*E-G*). Live imaging of third instar larval brains expressing ND-51^i^ in type II neuroblasts showed decreased division. Control neuroblasts divided at least once during live imaging in 40 mins, but ND-51 depleted neuroblasts did not show a division in the same duration (Fig. S1*B*). Collectively, these observations show that complex I is a crucial regulator of neuroblast proliferation.

We evaluated the impact of complex I depletion on differentiation in the type II neuroblast lineage. We quantified the number of distinct progenitor cells, mINPs, GMCs and neurons within the type II neuroblast lineage by immunostaining for specific markers. We found that the numbers of mINPs (Ase+, Dpn+) were decreased in ND-51^i^, ND-39^i^ and ND-19^i^ expressing lineages as compared to mCherry^i^ expressing controls (Fig. 2*A* and *B*). We assessed the proliferation of mINPs by co-staining with Dpn and pH3. Depletion of ND-51, ND-39, and ND-19 resulted in a decreased number of mINPs labelled with pH3 within the lineage compared to controls (Fig. S2*A* and *B*). This reduction in mINP proliferation corresponded to a decrease in GMC (Ase+ Dpn-) numbers (Fig. 2*A* and *C*), subsequently leading to a diminished production of neurons (high nuclear Pros) (Fig. 2*D* and *E*). We evaluated whether apoptosis led to a decrease in cells in the type II neuroblast lineage by immunostaining the brains with cleaved Caspase 3. We did not find any discernible difference between control and complex I depleted neuroblast lineages with cleaved Caspase 3 and found that apoptosis was not a cause of decrease in lineage cells (Fig. S2*A*). To confirm the specificity of RNAi-mediated complex I depletion, we utilized an independent shRNA targeting ND-51^i^ (BL29534). ND-51^i^ (BL29534) expression with *pnt*-Gal4 also showed a significant decrease in mINPs, GMCs, and neurons within the lineage (Fig. S2*B-F*).

**Fig. 2.**
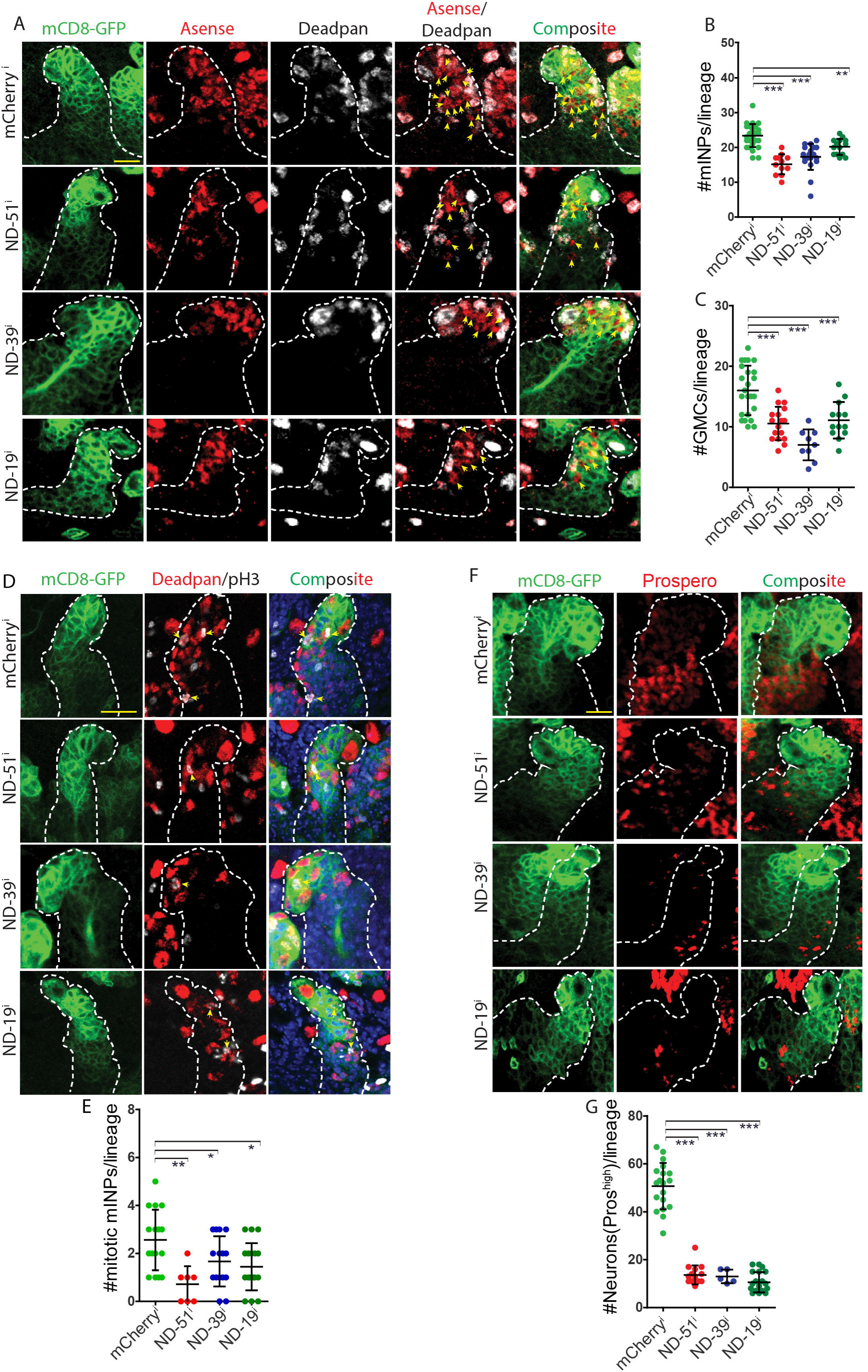
Complex I depletion in type II neuroblasts results in decreased mINPs, GMCs and neurons in each lineage. **(A)** Representative images of differentiated progenies in the type II neuroblast lineage of the mentioned genotypes. Dpn positive cells are mINPs (Gray). Dpn negative/Ase (red) positive cells are GMCs (Yellow arrowheads). Scale bar-10µm. **(B)** Quantification of mINPs in type II neuroblast lineages of genotypes, mCherry^i^ (n=28 type II neuroblast lineages, 8 brains), ND-51^i^ (12, 6), ND-391^i^ (20, 11), and ND-19^i^ (14, 9). **(C)** Quantification of GMCs in type II neuroblast lineages of genotypes, mCherry^i^ (n=24 type II neuroblast lineages, 11 brains), ND-51^i^ (17, 9), ND-391^i^ (9, 5), and ND-19^i^ (13, 9). **(D)** Representative images of immunostaining of neurons in type II neuroblast lineages of indicated genotypes. High nuclear Pros positive cells are neurons (red). Scale bar-10µm. **(E)** Quantification of neurons in type II neuroblast lineages of genotypes, mCherry^i^ (n=19 type II neuroblast lineages, 5 brains), ND-51^i^ (15, 3), ND-391^i^ (5, 2), and ND-19^i^ (20, 4).

Complex I depletion has also been shown to decrease the proliferation of tumor neuroblasts (7). In this analysis, we further find that the reduction in neuroblast and mINP proliferation resulting from complex I depletion in type II neuroblast lineage led to a decrease in the proliferation of transit amplifying cells and differentiated cells.

### Expression of Drp1 mutant in complex I deficient type II neuroblasts alleviates their proliferation and differentiation defect

Our prior analysis demonstrated that depleting Opa1 in type II neuroblasts leads to mitochondrial fragmentation and differentiation defects (6). Similar observations of mitochondrial fragmentation and impairment of differentiation have been reported in *Drosophila* intestinal stem cells upon Opa1 depletion (18). Drp1 activity is essential for driving mitochondrial fragmentation in Opa1 depleted cells (18–20). Additionally, concurrent depletion of Drp1 with Opa1 restores mitochondrial fusion and alleviates the differentiation defects, underscoring the necessity of a fused mitochondrial morphology for proper stem cell differentiation (6, 18). Depletion of ETC complex I and complex V in type I neuroblasts causes mitochondrial fragmentation (7). We assessed mitochondrial morphology in ND-51 depleted type II neuroblasts and muscle cells by crossing the shRNA lines with *pnt*-Gal4 and *mhc*-Gal4, respectively. As expected, ND-51 depleted type II neuroblasts contained fragmented mitochondria surrounding the nucleus and relatively smaller mitochondria in muscle cells (Fig. 3*A*).

**Fig. 3.**
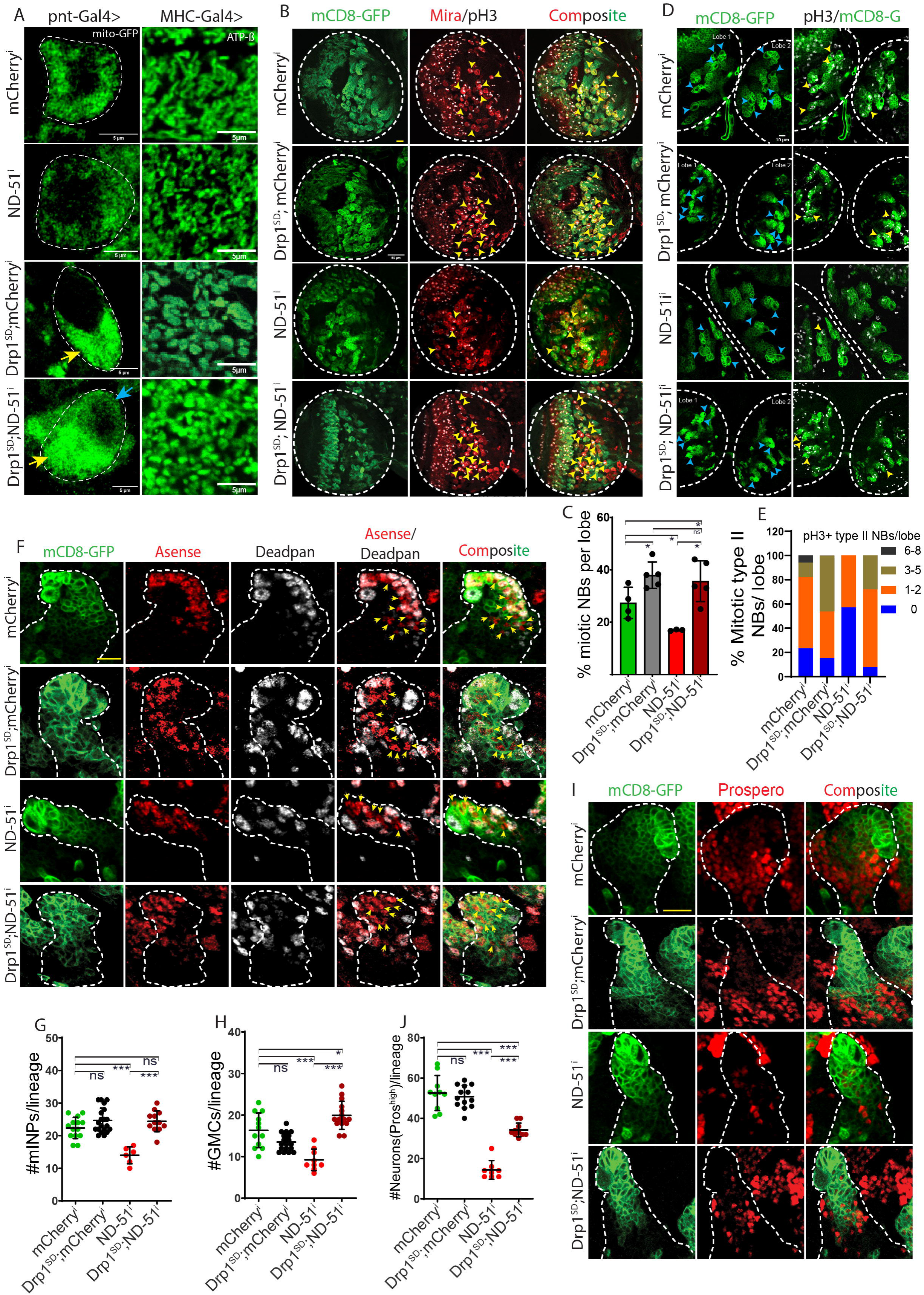
Mitochondrial fusion by Drp1^SD^ expression in complex I depleted neuroblasts alleviates the proliferation and differentiation defects. **(A)** Representative images of mitochondria stained with ATPb in type II neuroblasts and third instar larval muscle of the indicated genotypes. ND-51^i^ induces mitochondrial fragmentation in neuroblasts and muscle. Overexpression of Drp1^SD^ in ND-51^i^ neuroblasts results in clustered mitochondria. Drp1^SD^;mCherry^i^ neuroblasts show clustered mitochondria localized to one side (Yellow arrowhead), Drp1^SD^;ND-51^i^ neuroblasts exhibit clustered (yellow arrowhead) and fewer punctuated mitochondrial populations (blue arrowhead). **(B)** Representative images of pH3 labelled NSCs (Yellow arrowheads) in *Drosophila* third instar larval brain lobe of indicated genotypes. *wor*-Gal4 driven UAS-mCD8-GFP marks the boundary of stem cells and its lineage (green). Miranda labels stem cell cortex (red) and pH3 (grey). Scale bar-10µm. **(C)** Graph showing the quantification of percent mitotically dividing NSCs present in each brain lobe of third instar larvae, expressing *wor*-Gal4 driven mCherry^i^ (n=295 NSCs, 3 brains), Drp1^SD^; mCherry^i^ (476, 5), ND-51^i^ (267, 3), Drp1^SD^; ND-51^i^ (488, 5). **(D)** Representative images of pH3 positive type II neuroblasts (Yellow arrowheads) in *Drosophila* third instar larval brain of indicated genotypes. *pnt*-Gal4 driven mCD8-GFP marks the boundary of stem cells (Blue arrowheads) and its lineage (green), and pH3 (grey). Scale bar-10µm. **(E)** Quantification of the number of pH3 positive type II neuroblasts in each brain lobe of the larval brain of genotype, mCherry^i^ (n=144 NSCs, 9 brains), Drp1^SD^; mCherry^i^ (112, 7), ND-51^i^ (160, 10), Drp1^SD^; ND-51^i^ (200, 13). **(F)** Representative images of immunostaining of mINPs and GMCs cells in type II neuroblast lineage of indicated genotypes. Dpn (Gray), Ase (red)/Dpn negative cells are GMCs (Yellow arrowheads). Scale bar-10µm. **(G)** Quantification of mINPs in type II neuroblast lineages of genotypes, mCherry^i^ (n=28 type II neuroblast lineages, 11 brains), Drp1^SD^; mCherry^i^ (20, 7), ND-51^i^ (12, 9), Drp1^SD^; ND-51^i^ (12, 6). **(H)** Quantification of GMCs in type II neuroblast lineages of genotypes, mCherry^i^ (n=12 type II neuroblast lineages, 3 brains), Drp1^SD^; mCherry^i^ (18, 5), ND-51^i^ (14, 3), Drp1^SD^; ND-51^i^ (15, 5). **(I)** Representative images of neurons labelled with high nuclear Pros (red) immunostaining in type II neuroblast lineages of indicated genotypes. Scale bar-10µm. **(J)** Quantification of neurons in type II neuroblast lineages of genotypes mCherry^i^ (n=10 type II neuroblast lineages, 5 brains), Drp1^SD^; mCherry^i^ (13, 5), ND-51^i^ (8, 5), Drp1^SD^; ND-51^i^ (10, 3).

We investigated whether Drp1 depletion by co-expression of the dominant negative form of Drp1 (Drp1^SD^) with complex I RNAi could reverse the mitochondrial morphology defects. For this, we generated the combinations, Drp1^SD^;ND-51^i^ and Drp1^SD^;mCherry^i^ as a control and crossed them to *pnt*-Gal4 and *mhc*-Gal4 to assess mitochondrial morphology. Mitochondria in Drp1^SD^;mCherry^i^ type II neuroblasts were clustered and localised to one side of the cell as compared to mCherry^i^ alone.

Drp1^SD^;mCherry^i^ expression in muscle cells led to bigger and interconnected mitochondria as compared to mCherry^i^ alone. Drp1^SD^;ND-51^i^ neuroblasts contained a mixed population of clustered mitochondria localised to one side of the cell and fragmented mitochondria. Muscle cells expressing Drp1^SD^;ND-51^i^ showed larger mitochondria than ND-51^i^ alone.

We further assessed the proliferation of Drp1^SD^;ND-51^i^ expressing neuroblasts using *wor*-gal4 and *pnt*-Gal4. In each brain lobe, the proportion of mitotically dividing neuroblasts was significantly increased in Drp1^SD^;ND-51^i^ compared to ND-51^i^ alone (Fig. 3*B*-*E*). Next, we analysed the consequence of Drp1 depletion on differentiation in the ND-51^i^ expressing type II neuroblast lineage.

Consistent with the suppression of the defect in proliferation of neuroblasts (Fig. 3*A*-*E*), the number of mINPs increased in Drp1^SD^;ND-51^i^ compared to ND-51^i^ alone (Fig. 3*F* and *G*). Furthermore, we observed an increase in mitotically dividing mINPs in Drp1^SD^;ND-51^i^ compared to ND-51^i^ (Fig. S3*A* and *B*). This increased division of mINPs in Drp1^SD^;ND-51^i^ type II neuroblast lineage resulted in the rescue of the number of GMCs (Fig. 3*F-H*), and the neurons were also partially rescued (Fig. 3*I* and *J*). To confirm that the rescue of differentiation defects upon Drp1^SD^ expression was not a consequence of Gal4 dilution, we expressed UAS-mito-GFP with ND-51^i^ and found that the numbers of mINPs, GMCs, and neurons were comparable to ND-51^i^ alone (Fig. S2*B-F*). In summary, mitochondrial fusion suppresses the differentiation defects caused by complex I depletion in the type II NB lineage.

### Depletion of Drp1 suppresses the defects in G1/S progression and cyclin E nuclear accumulation in complex I knockdown type II neuroblasts

Mitochondrial activity and dynamics regulate cell cycle progression through the G1/S phase. Depletion of complex I in *Drosophila* eye discs causes a delay in the G1/S transition and decrease in cyclin E, mediated by high ROS levels and JNK-pathway activity (21, 22). Drp1 depletion also affects cyclin E levels in *Drosophila* follicle cells (23, 24). We assessed the impact of complex I depletion on G1-S progression and cyclin E accumulation in the nucleus. We incubated the live brains with EdU (5-ethynyl-2’-deoxyuridine), a thymidine nucleoside analog that incorporates into newly synthesised DNA during the S-phase of the cell cycle. ND-51^i^ expressing type II neuroblast lineage showed a decreased number of EdU-positive cells and mINPs compared to controls (Fig. 4*A*-*C*). This decrease was restored in Drp1^SD^;ND-51^i^ expressing type II neuroblasts and their lineage. There was a decrease in the nuclear accumulation of cyclin E in ND-51^i^ expressing neuroblasts compared to the controls (Fig. 4*D* and *E*). Further, cyclin E levels in Drp1^SD^;ND-51^i^ expressing neuroblasts were comparable to control and greater than ND-51^i^ alone (Fig. 4*D* and *E*). Therefore, complex I disruption induces G1/S phase transition defects by decreasing nuclear cyclin E in type II neuroblasts.

**Fig. 4.**
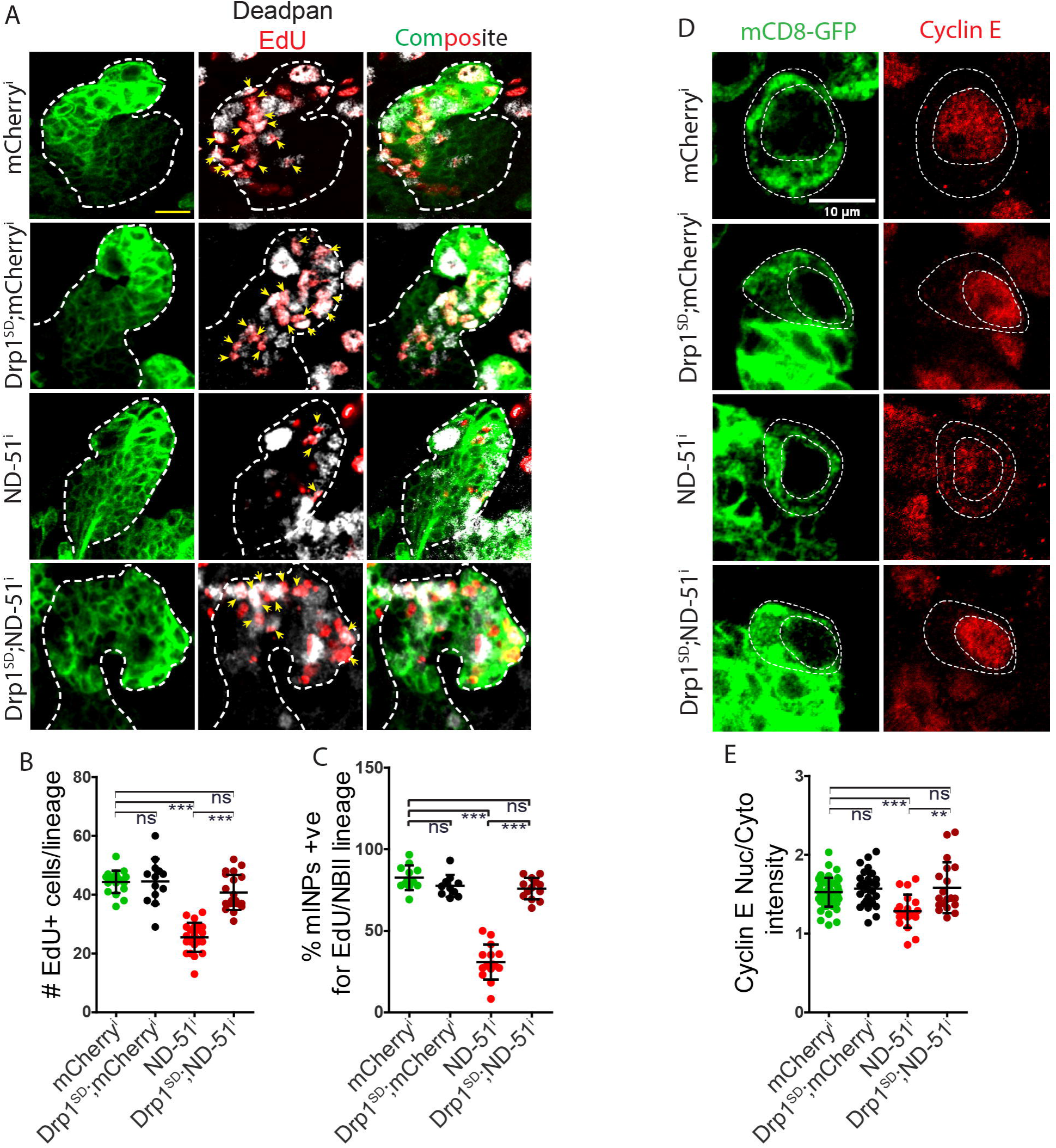
Depletion of complex I in type II neuroblasts inhibits G1-S phase progression in mINPs by reducing Cyclin E levels and this defect is rescued by Drp1^SD^ expression. **(A)** Representative images of cyclin E immunostaining (red) in type II neuroblasts (encircled by a dashed line) of indicated genotypes. Scale bar-10µm. **(B)** Quantification of cyclin E intensity in nucleus with respect to cytoplasm of type II neuroblasts of following genotypes: mCherry^i^ (n=62 type II neuroblasts, 9 brains), Drp1^SD^;mCherry^i^ (28, 3), ND-51^i^ (20, 3), Drp1^SD^;ND-51^i^ (19, 3). **(C)** Representative images of EdU positive cells (red) in the lineages of type II neuroblast of mentioned genotypes. Scale bar-10µm. **(D)** Quantification of percent EdU positive mINPs (Yellow arrowheads) in the lineages of type II neuroblasts of genotypes:mCherry^i^ (n=12 type II neuroblasts lineages, 6 brains), Drp1^SD^;mCherry^i^ (11, 5), ND-51^i^ (15, 6), Drp1^SD^;ND-51^i^ (13, 4). **(E)** Quantification of total EdU positive cells in the lineages of type II neuroblasts of genotypes:mCherry^i^ (n=19 type II neuroblasts lineages, 6 brains), Drp1^SD^;mCherry^i^ (13, 5), ND-51^i^ (23, 6), Drp1^SD^;ND-51^i^ (20, 4).

### Decreased MMP, mitochondrial fragmentation, decreased cristae numbers and increased ROS production in complex I depleted type II neuroblasts is rescued by mitochondrial fusion

Disruption of the mitochondrial electron transport chain (ETC) in neuroblasts leads to mitochondrial fragmentation, decreased mitochondrial activity, and altered metabolites (8, 25). We measured the changes in MMP, cristae organization, and ROS generation to assess mitochondrial functionality in complex I depleted type II neuroblasts. We used the potentiometric dye, Tetra-methyl-rhodamine methyl ester (TMRM), to assess MMP in living third-instar larval brains. TMRM fluorescence was measured in type II neuroblasts and expressed as a ratio to neighbouring control type I neuroblasts. We observed a significant reduction in MMP in ND-51 and ND-19 depleted type II neuroblasts (Fig. 5*A* and *B*). Mitochondrial fusion by Drp1^SD^ expression did not alter the MMP, and co-expression of Drp1^SD^ with ND-51^i^ and ND-19^i^ in type II neuroblasts restored the MMP.

**Fig. 5.**
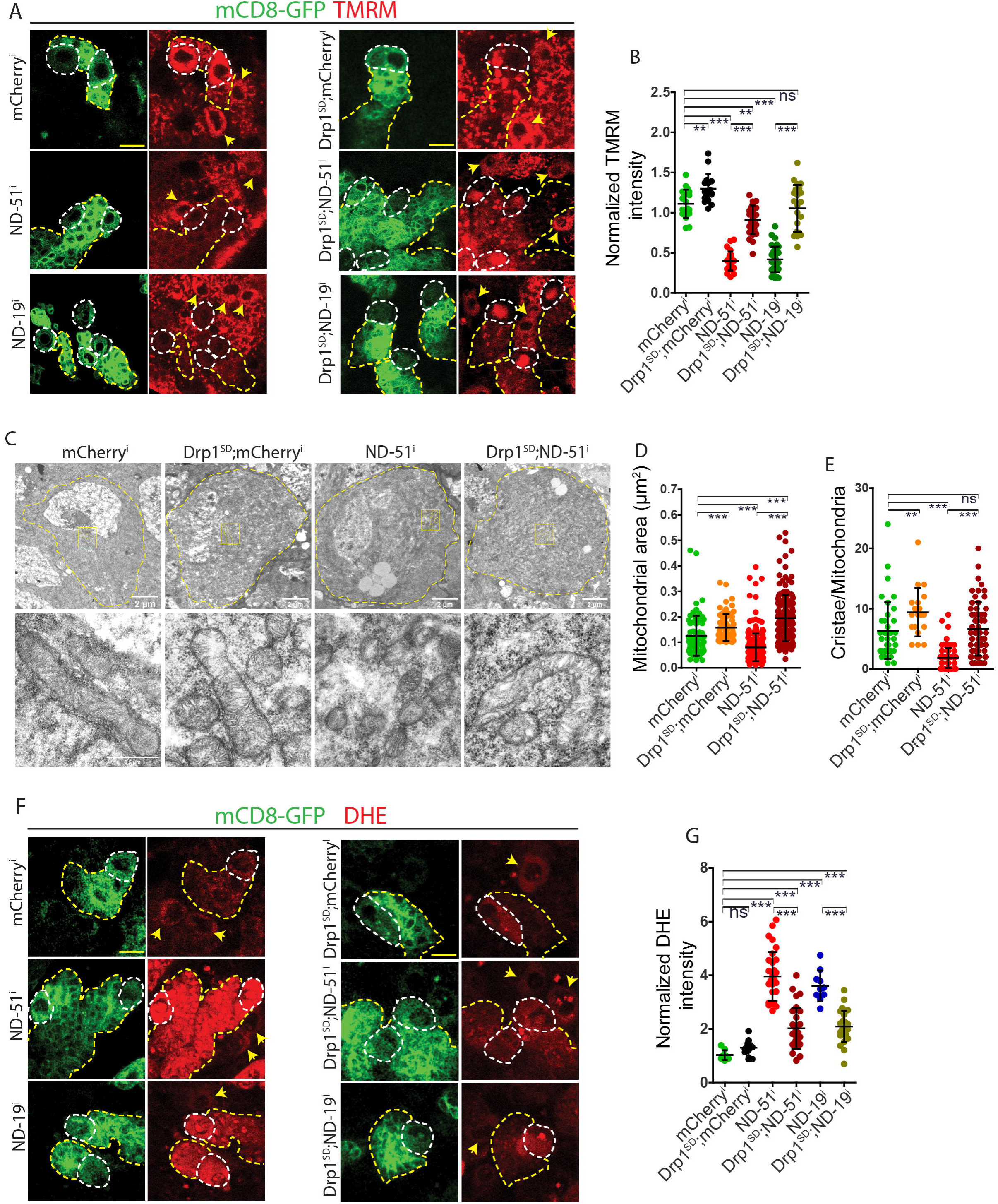
Depletion of complex I decreases MMP, cristae and ROS in type II neuroblasts and mitochondrial fusion alleviates these defects. **(A)** Representative images of TMRM fluorescence in type II neuroblasts (red, circled by white dashed line). Surrounding control type I neuroblasts are shown by yellow arrow heads. Scale bar-10µm. **(B)** Quantification of TMRM intensity in type II neuroblast, normalized to the intensity of neighbouring type I neuroblast of the genotypes: mCherry^i^ (n=19 type II neuroblasts, 5 brains), Drp1^SD^;mCherry^i^ (16, 5), ND-51^i^ (17, 9), Drp1^SD^;ND-51^i^ (28, 5), ND-19^i^ (34, 6), Drp1^SD^;ND-19^i^ (22, 4). **(C)** Representative electron micrographs of mCherry^i^ neuroblasts (elongated and well-developed cristae mitochondria), Drp1^SD^;mCherry^i^ (fused, cristae-rich mitochondria clustered on one side of the neuroblasts), complex I depletion (ND-51^i^) (fragmented and circular mitochondria) and Drp1^SD^;ND-51^i^ (mitochondria display a mixed morphology of fragmented and fused forms, predominantly clustered on one side with cristae). **(D)** Quantification of mitochondrial area in the following genotypes: mCherry^i^ (n=76 mitochondria, 10 neuroblasts, 3 brains), Drp1^SD^;mCherry^i^ (66, 8, 3), ND-51^i^ (247, 12, 3), Drp1^SD^;ND-51^i^ (138, 16, 3). **(E)** Quantification of cristae number in following genotypes: mCherry^i^ (n=39 mitochondria, 10 neuroblasts, 3 brains), Drp1^SD^;mCherry^i^ (20, 8, 3), ND-51^i^ (85 mitochondria, 12, 3), Drp1^SD^;ND-51^i^ (60, 16, 3). **(F)** Representative images showing DHE fluorescence (red, encircled by white dashed line). Control type I neuroblasts are marked by yellow arrow heads. Scale bar-10µm. **(G)** Quantification of DHE fluorescence intensity in type II neuroblast normalized to surrounding type I neuroblast of following genotypes: mCherry^i^ (n=9 type II neuroblasts, 5 brains), Drp1^SD^;mCherry^i^ (11, 5), ND-51^i^ (32, 7), Drp1^SD^;ND-51^i^ (30, 5), ND-19^i^ (10, 3), Drp1^SD^;ND-19^i^ (26, 5).

The inner mitochondrial membrane in fused mitochondria can accommodate well-organised lamellar cristae, whereas fragmented mitochondria contain poor cristae morphology. From a functional perspective, cristae expand the surface area of the inner mitochondrial membrane, thereby increasing ETC components, for increased oxidative phosphorylation leading to higher MMP (26). By concentrating proteins involved in oxidative phosphorylation and lowering the mean distance between electron transport chain (ETC) complexes, cristae behave as specialised compartments that ensure favourable conditions for ATP synthesis (27–29). Therefore, cristae disruption might be the reason for reduced MMP on complex I depletion in neuroblasts. To study cristae organization, we used transmission electron microscopy on brains expressing ND-51^i^ and mCherry^i^ with *wor*-Gal4. Control neuroblasts have relatively elongated mitochondria with cristae-rich morphology. ND-51^i^ neuroblasts possess fragmented mitochondria; the mitochondria are smaller and have less crossectional area. The ND-51^i^ neuroblasts have fewer numbers and poorly organised cristae than controls (Fig. 5*C-E*). In Drp1^SD^;ND-51^i^ expressing type II neuroblasts, mitochondrial morphology and cristae number were similar to Drp1^SD^;mCherry^i^ controls (Fig. 5*C-E*). These observations show that the rescue of MMP by mitochondrial fusion in complex I depleted neuroblasts is coincident with the achievement of appropriate mitochondrial morphology and cristae architecture.

In *Drosophila* embryos, inhibiting ETC activity induces cellular ATP stress and increased phosphorylated AMP-activated protein kinase (pAMPK) (19). Low ATP levels elevate AMP, leading to increased pAMPK (30). Incubating third-instar larval brains with a non-hydrolysable glucose analogue 2-deoxyglucose also increases pAMPK (6). To investigate if complex I depletion decreased ATP in type II neuroblasts, we assessed ATP levels in ND-51 depleted type II neuroblasts using the BioTracker ATP-Red Live Cell Dye. The fluorescence of the ATP-Red dye was found to be comparable in complex I depleted and control type II neuroblasts (Fig. S4*A* and *B*). Additionally, immunostaining for pAMPK in ND-51^i^ expressing type II neuroblasts gave fluorescence similar to controls, suggesting that ATP levels are similar to controls in these cells (Fig. S4*C* and *D*).

Alterations in mitochondrial morphology and ETC change ROS levels in the mitochondria and the cytoplasm (18, 31, 32). Complex I of the ETC generates ROS, and ROS are recognised as crucial signalling molecules in cellular processes, including stem cell maintenance and differentiation (18, 33–36). Superoxides generated in the mitochondria can diffuse into the cytoplasm as H_2_O_2_ and also be transported outside via the voltage gated anion channels (VDAC) in the outer membrane (37). We assessed the ROS levels in type II neuroblasts using the fluorescent dye dihydroxy ethidium (DHE). DHE fluorescence in type II neuroblasts was estimated as a ratio to neighbouring control type I neuroblasts. Knockdown of complex I subunits ND-51 and ND-19 in type II neuroblasts resulted in a significant elevation in ROS levels as compared to controls and the ROS levels were restored in both Drp1^SD^;ND-51^i^ and Drp1^SD^;ND-19^i^ expressing neuroblasts (Fig. 5*F* and *G*).

These data showed that fusion of mitochondria by Drp1^SD^ expression in complex I depleted neuroblasts restored the mitochondrial cristae architecture and mitochondrial activity defects as measured by MMP and ROS levels.

### Depletion of ROS scavenging enzymes leads to a decrease in type II neuroblast differentiation and G1-S transition

To verify whether ROS regulates type II neuroblast differentiation, we manipulated ROS levels in type II neuroblasts by knocking down the ROS scavenger enzymes, the cytoplasmic superoxide dismutase SOD1 and the mitochondrial superoxide dismutase SOD2. We downregulated SOD1 and SOD2 by expressing RNAi.

Depleting SOD enzymes in type II neuroblasts and larval muscle showed mitochondrial fragmentation (Fig. S5E), but interestingly, MMP was not altered in SOD1^i^ type II neuroblasts but decreased slightly in SOD2^i^ (Fig. S5F and G). SOD1/2 downregulation in type II neuroblasts resulted in elevated ROS levels (Fig. S5*A* and *B*) and decreased number of mINPs, GMCs, and neurons in the lineages as compared to controls (Fig. 6*A-E*). Downregulation of SOD1/2 also led to a decreased number of total EdU positive cells in the lineage, and the proportion of mINPs positive for EdU also declined (Fig. 6*F-H*). This decreased proliferation of cells in type II neuroblast lineage was consistent with the decrease of nuclear cyclin E localisation (Fig. 6*I* and *J*). These results show that enhanced ROS generation in type II neuroblasts impedes nuclear cyclin E levels, leading to proliferation defects and ultimately resulting in a decreased number of differentiated cells in the lineage. ROS is therefore causally linked to proliferation and defects caused by complex I depletion in type II neuroblasts.

**Fig. 6.**
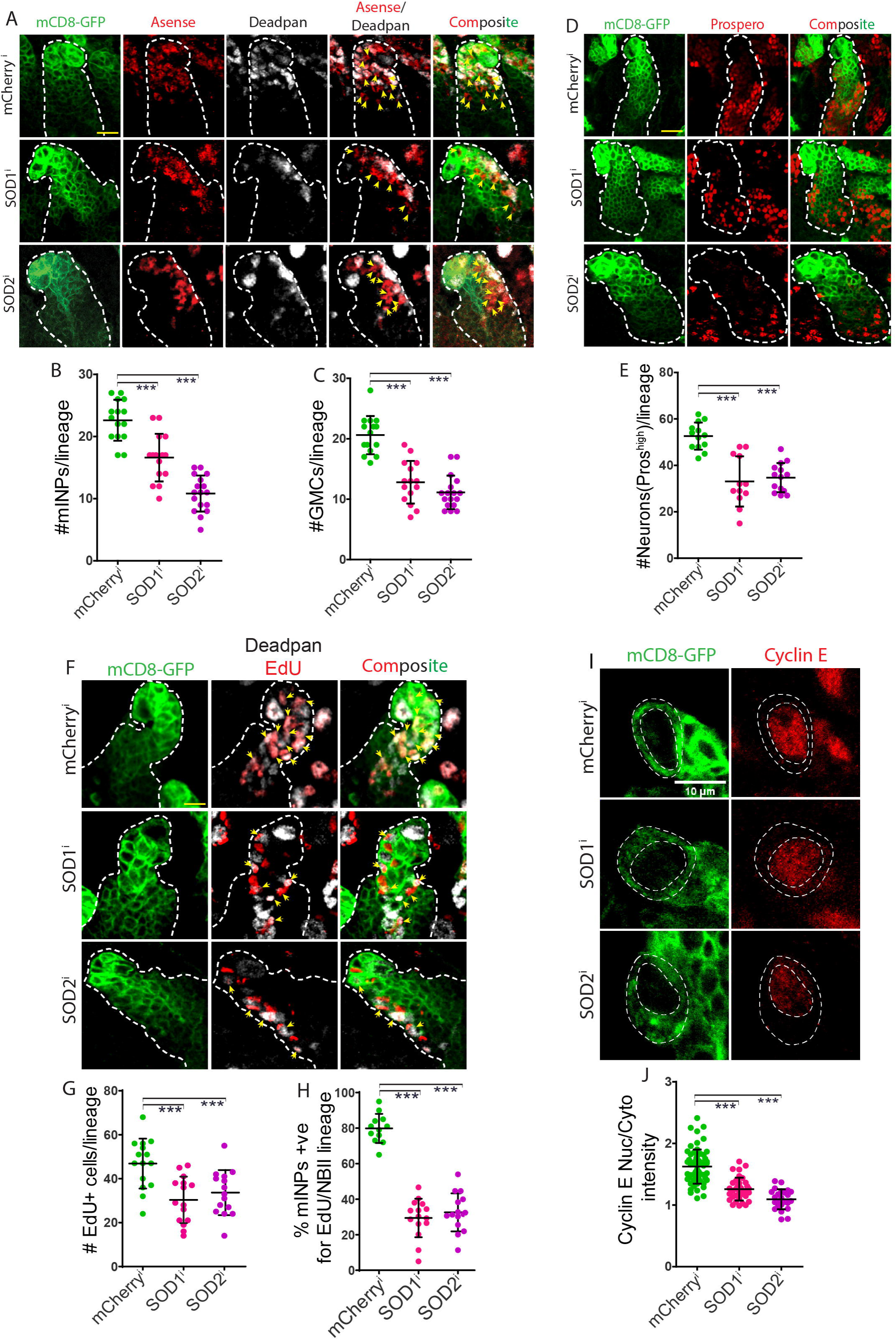
Depletion of ROS scavenging enzymes leads to decreased mINPs, GMCs, and neurons in each type II neuroblast lineage due to a defect in G1-S transition. **(A)** Representative image showing mINPs and GMCs of mentioned genotypes. Dpn positive mINPs (Gray). Ase positive (red)/Dpn negative GMCs (Yellow arrowheads). **(B)** Quantification of mINPs in type II neuroblast lineages of genotype: mCherry^i^ (n=15 type II neuroblast lineages, 3 brains), SOD1^i^ (15, 4), and SOD2^i^ (17, 4). **(C)** Quantification of GMCs in type II neuroblast lineages of genotypes: mCherry^i^ (n=15 type II neuroblast lineages, 3 brains), SOD1^i^ (15, 4), and SOD2^i^ (17, 4). **(D)** Representative images of neurons labelled with high nuclear Pros (red) immunostaining in type II neuroblast lineages of indicated genotypes. Scale bar-10µm. **(E)** Quantification of neurons in type II neuroblast lineages of genotypes: mCherry^i^ (n=13 type II neuroblast lineages, 3 brains), SOD1^i^ (12, 6), and SOD2^i^ (15, 4). **(F)** Representative images of EdU positive cells (red) in type II neuroblasts lineages of mentioned genotypes. Scale bar-10µm. **(G)** Quantification of percent EdU positive mINPs (Yellow arrowheads) in type II neuroblast lineages of genotypes: mCherry^i^ (n=12 type II neuroblast lineages, 5 brains), SOD1^i^ (15, 5), and SOD2^i^ (15, 3). **(H)** Quantification of total EdU positive cells (red) in type II neuroblast lineages of genotypes:mCherry^i^ (n=15 type II neuroblast lineages, 5 brains), SOD1^i^ (15, 5), and SOD2^i^ (15, 3). **(I)** Representative images of cyclin E immunostaining (red) in type II neuroblasts of indicated genotypes. Type II neuroblasts are shown as encircled by white dashed line. Scale bar-10µm. **(J)** Quantification of cyclin E intensity in nucleus with respect to cytoplasm of type II neuroblast of following genotypes: mCherry^i^ (n=64 type II neuroblasts, 9 brains), SOD1^i^ (12, 4), and SOD2^i^ (24, 4).

### Overexpression of ROS scavengers in complex I depleted type II neuroblast suppresses differentiation and G1/S transition defects

Since elevated ROS via complex I depletion or depletion of superoxide scavenging enzymes led to loss of differentiation in the type II neuroblasts, we assessed if decreasing the ROS levels in ND-51^i^ by overexpression of SOD2 would alleviate the differentiation defect in ND-51^i^. Overexpression of SOD2 in type II neuroblasts did not change ROS levels (Fig. S5*C* and *D*) and the number of mINPs, GMCs and neurons as compared to controls. But overexpression of SOD2 in ND-51 depleted type II neuroblasts (SOD2^OE^;ND-51^i^) decreased ROS levels compared to ND-51^i^ alone (Fig. S5*C* and *D*). Further, the SOD2^OE^;ND-51^i^ combination showed a restoration of the numbers of mINPs, GMCs and neurons in each type II neuroblast lineage as compared to ND-51^i^ alone (Fig. 7*A-E*). Rescue of differentiation defects in SOD2^OE^;ND-51^i^ type II neuroblasts was consistent with the rescue of S-phase proliferation defects in the lineage, the number of EdU positive cells and proportion of EdU positive mINPs were higher than the ND-51^i^ alone (Fig. S7*A-C*). Also, nuclear cyclin E levels in SOD2^OE^;ND-51^i^ type II neuroblasts were similar to those observed in controls (Fig. S6*D* and *E*). SOD2^OE^;ND-51^i^ expressing type II showed a loss of MMP as compared to controls (Fig. S5*F* and *G*) and this was similar to like ND51^i^ (Fig. 5*A* and *B*), suggesting that proliferation and differentiation defects on complex I depletion are because of high ROS but not MMP loss. Since mitochondrial fusion by Drp1 depletion in complex I downregulated type II neuroblast rescued differentiation defects (Fig. 3) by lowering ROS levels (Fig. 5*F* and *G*), we expressed Drp1^SG^ in SOD2^i^ type II neuroblast. We observed that SOD2^i^ mediated differentiation defects were rescued; the number of mINPs, GMCs and neurons were increased compared to SOD2^i^ alone and were non-significantly different from control (Fig. S7*A-E*). These findings show a crucial role for maintenance of appropriate ROS levels by mitochondrial fusion and complex I activity in regulating differentiation in the type II neuroblast lineage.

**Fig. 7.**
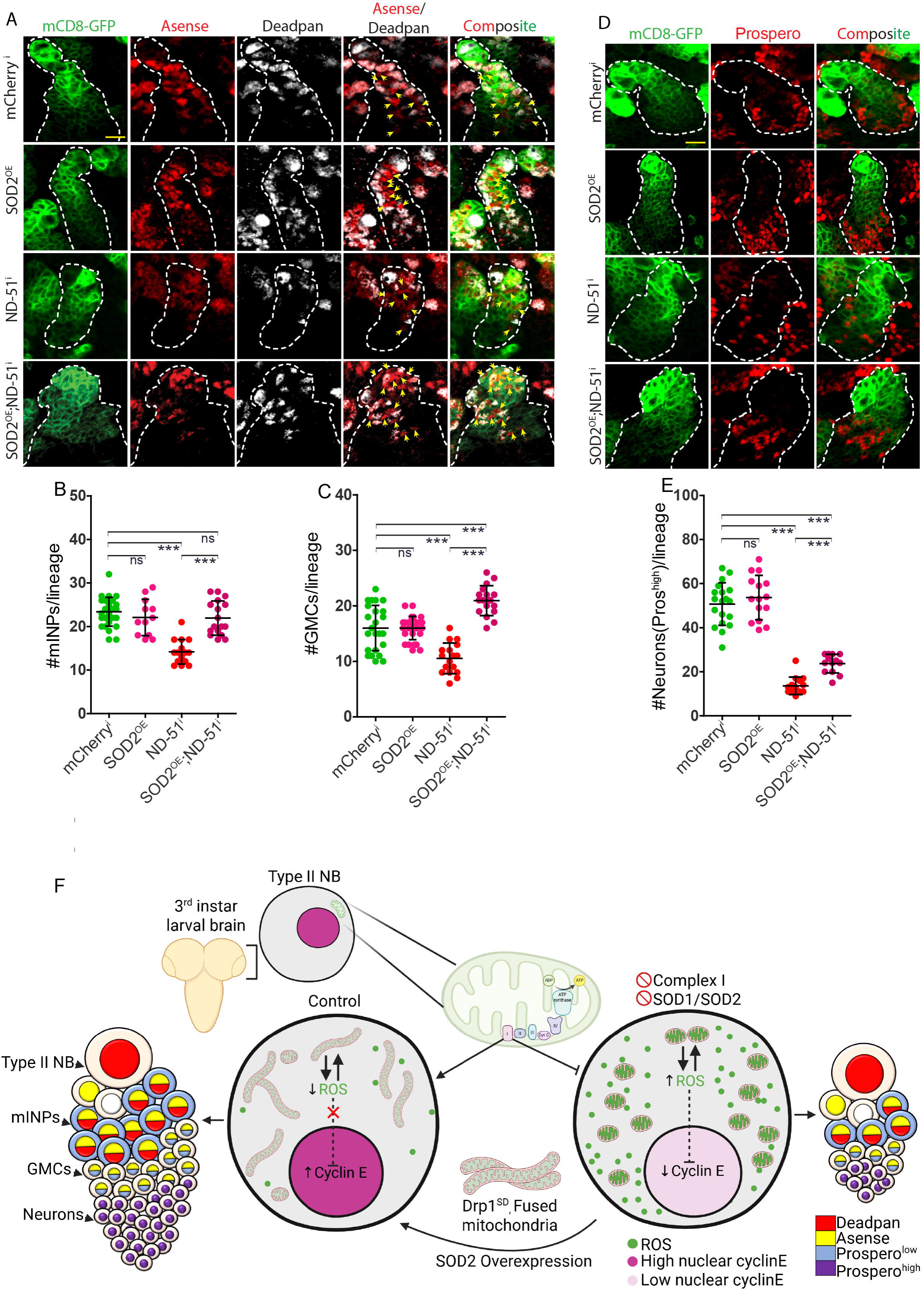
Overexpression of SOD2 in ND-51^i^ expressing type II neuroblast alleviates differentiation defects. **(A)** Representative image showing immunostaining of mINPs and GMCs if type II neuroblast lineage of mentioned genotypes. Dpn positive mINPs (Gray). Ase positive (red)/Dpn negative cells GMCs (Yellow arrowheads). **(B)** Quantification of mINPs in type II neuroblast lineages of genotype: mCherry^i^ (n=21 type II neuroblast lineages, 6 brains),SOD2^OE^ (12,3), ND-51^i^ (15,3), and SOD2^OE^;ND-51^i^ (17, 5). **(C)** Quantification of GMCs in type II neuroblast lineages of genotypes: mCherry^i^ (n=24 type II neuroblast lineages, 6 brains), SOD2^OE^ (19,3), ND-51^i^ (17,3), and SOD2^OE^;ND-51^i^ (18,5). **(D)** Representative images of neurons marked with high nuclear Pros (red) in type II neuroblast lineages of indicated genotypes. Scale bar-10µm. **(E)** Quantification of neurons in type II neuroblast lineages of genotypes: mCherry^i^ (n=19 type II neuroblast lineages, 5 brains), ND-51^i^ (15, 3), SOD2^OE^ (15,4), SOD2^OE^;ND-51^i^ (13, 4). **(F)** Schematic summary showing that complex I depletion impairs neuroblast proliferation and differentiation due to mitochondrial fragmentation and increased ROS leading to cell cycle defects. Mitochondrial fusion and SOD2 overexpression in complex I depleted neuroblasts leads to suppression of cell cycle defects.

In summary, the elevation of ROS levels in type II neuroblasts, induced by either complex I or SOD depletion, leads to mitochondrial fragmentation and differentiation defects by abrogating the G1/S phase transition. However, restoring ROS levels by overexpressing scavenging enzymes or fusing mitochondria by depleting Drp1 in these neuroblasts rescues the differentiation defects (Fig. 7*F*). Our studies demonstrate that relatively fused mitochondrial morphology plays a crucial role as an essential regulator of ETC activity and ROS, facilitating progression through the cell cycle for differentiation in neuroblasts of *Drosophila*.

## Discussion

We show that optimal complex I activity and mitochondrial fusion limits ROS to regulate the cell cycle and differentiation in the *Drosophila* type II neuroblast lineage. We discuss our results in the context of the following: 1] the mechanism by which there is a change in ROS by complex I depletion and how mitochondrial fusion impacts this change and 2] the mechanism of control of mitochondrial activity and morphology in *vivo* in stem cell differentiation.

During electron transport through ETC complexes, 0.2-2% of electrons escape ETC and interact with oxygen to form superoxides (38). Complex I and complex III, are the main sites of ROS production in mitochondria (39). Dysfunctionality of complex I or its inhibition by rotenone leads to inefficient transfer of electrons, leading to their build-up and increase in ROS by reaction with oxygen in the mitochondria (40). We observed an increase of ROS on complex I depletion by depleting its subunits ND-51 or mammalian NDUFV1 and ND-19 or mammalian NDUFA8. ND-51 belongs to the core hydrophilic matrix and catalyses the oxidation of NADH to NAD+ and is part of ∼815kDa complex I assembly intermediate.

Downregulation of ND-51 and ND-19 in *Drosophila* muscle decreases complex I assembly (13). Thus, disrupting complex I formation in type II neuroblasts on ND-51 and ND-19 knockdown may cause complex I to lose electrons and produce ROS. Reverse electron transfer (RET) may be another possibility for generation of increased ROS. RET catalyses the reduction of matrix NAD+ with electrons provided by CoQH2. The electrons from complex I and II are transferred to CoQH2, some are further moved to complex III (FET, forward electron transfer), and others flow back to complex I (RET). Excessive ROS is generated during RET compared to FET, presumably due to the dissociation of FMNH_2_ from complex I and the reaction with matrix O_2_ (41). FMN and the CoQ reduction sites of complex I are centres for ROS production during RET (42, 43).

In complex I deficient neuroblasts, mitochondrial fusion might reduce ROS generation via facilitating ETC supercomplex formation in the cristae. These supercomplexes increase electron transport efficiency between ETC complexes by optimising the distance between them (44, 45). Hydrogen peroxide, a small and electrically neutral molecule, diffuses through the inner and outer membranes of mitochondria to reach the cytoplasm (46). Superoxides on the other hand may pass to the cytoplasm through mitochondrial channels. Increased ROS accumulation beyond a threshold level in mitochondria induces the opening of mitochondrial channels, mitochondrial permeability transition (MPT) pore (47) and inner membrane anion channel (IMAC) (48), which releases ROS to the cytoplasm, accompanied by MMP dissipation (49). ROS accumulation in the cytosol promotes the opening of MPT and IMAC in other mitochondria, called ROS-induced ROS release (RIRR). Therefore, RIRR might be another reason for MMP loss on complex I depletion and mitochondrial fusion rescuing MMP by inhibiting RIRR through decreasing ROS levels.

Decrease in complex I subunits leads to mitochondrial fragmentation. This is likely to occur due to the alteration of mitochondrial fusion and fission proteins or the impact of ROS on mitochondrial fusion and fission proteins. Complex I inhibition by rotenone induces mitochondrial fission in a Drp1 dependent manner in neurons in the Parkinson’s disease model (50). Decreasing mitochondrial fission by expressing a dominant-negative Drp1 or overexpressing Mfn led to the inhibition of apoptosis of these neurons. Increase in ROS has also been shown to increase expression of Drp1 and activate Drp1 fission by increasing phosphorylation at the 616 residue in mouse hippocampal neurons (32, 51). ROS can regulate mitochondrial dynamics by modulating fusion and fission proteins at the transcriptional level and through post-translational modifications. High cellular ROS activates MAP kinases like ERK, which then phosphorylate and activate Drp1, promoting fission (52). In hepatocellular carcinoma, elevated ROS generation facilitates survival by activating the AKT pathway, which then activates NF-κB. This pathway leads to the expression of proteins involved in mitochondrial fission, like Drp1, and suppresses fusion proteins MFN1 (53).

Various signalling pathways regulate mitochondrial activity and stem cell differentiation. Notch signalling regulates mitochondrial morphology and thereby affects stem cell fate decisions. In mice, less Notch signaling maintains fragmented mitochondria and promotes intestinal stem cells and neural stem cells differentiation (33, 54). Fused mitochondria morphology promotes the differentiation of *Drosophila* intestinal stem cells by maintaining low ROS and suppressing JNK pathway. Inducing mitochondrial fragmentation elevated ROS levels and JNK activity, and inhibited differentiation (18). This study motivates an analysis of the signaling pathways that regulate mitochondrial activity and morphology in type II neuroblast differentiation and the various metabolites that need to be restricted by mitochondrial fusion to promote neural stem cell differentiation.

## Acknowledgements

We thank the RR lab for continuous feedback on this study. We thank the IISER Pune *Drosophila* and microscopy facility for help with stocks and microscopy. We thank Flybase and Bloomington Stock center for help with fly stocks. We thank Jurgen Knoblich (IMB, Vienna, Austria) for stocks. We thank Yuh-Nung Jan, UCSF, USA for Asense antibody.

## Material and methods

### Genetic crosses and fly lines

10 female flies were placed along with 5-6 male flies in polystyrene vials containing cornmeal agar media at 29°C. After 48 hours, the flies were transferred to fresh vials. 5-6 days after, third instar larvae emerged, and subsequent analysis was performed. The fly lines used in the study are: *wor*-Gal4, *pnt*-Gal4, mCD8-GFP (Jurgen Knoblich, IMP, Vienna, Austria), *mhc*-Gal4, ND-51^i^ (Bloomington stock, BL36701, BL29534), ND-39^i^ (BL 52922), ND-19^i^ (BL50577), Drp1^S193D^ (Drp1^SD^) (6, 55), hSOD1A4V (BL33607), mCherry^i^ (BL35785), mito-GFP (BL8442), SOD1^i^ (BL24750), SOD2^i^(BL25969), and SOD2^OE^ (BL24494)(56–58). The following combinations of transgenes were made by using standard fly genetics and appropriate balancers to bring the lines together: Drp1^SD^/CyOGFP;mCherry^i^/TM6B, Drp1^SD^/CyOGFP;ND-51^i^(36701)/TM6B, Drp1^SD^/CyOGFP;ND-19^i^ (50577)/TM6B, Drp1^SD^/CyOGFP;*pnt*-Gal4,CD8-GFP/TM6B, mitoGFP^OE^/CyOGFP;*pnt*-Gal4/TM6B, SOD2^OE^/CyOGFP;ND-51^i^(BL36701)/TM6B, Drp1^SG^/FM7a;SOD2^i^/TM3,SerGFP The *wor*-Gal4, UAS-mCD8-GFP was employed to express transgenes in all third-instar larval neuroblasts and *pnt*-Gal4, UAS-mCD8-GFP was utilised specifically for type II neuroblasts.

### Larval brain Immunofluorescence

Wandering third instar larvae were collected and the brain was dissected in Schneider’s media (Invitrogen-Thermofisher, 21720024), followed by 4% paraformaldehyde (Sigma, 158127) fixation for 25 minutes. The brains were then washed in 0.1% PBST (1X PBS, 0.1% Triton X-100 (Sigma,T8787)) for 30 minutes at room temperature (RT) and blocked with 1% BSA (Himedia, MB083) in 0.1% PBST for 1 hour at RT. Following this, the brains were incubated with primary antibodies (diluted in 1% BSA) overnight at 4D on a shaker. The brains were washed thrice (20min, 10min, 10min) with 0.1% PBST and incubated with fluorescently coupled secondary antibodies (diluted in 0.1% PBST) for 1 hour at RT followed by 20 minutes washing with 0.1% PBST. The brains were stained with the DNA dye Hoechst for 10 minutes, washed for 10 minutes with 0.1% PBST, and then mounted in SlowFade Gold Antifade Mountant (Invitrogen, S36937). The following dilutions of antibodies were used: chicken anti-GFP (1:1000, Invitrogen, A10262), rabbit anti-Ase (1:10000, Yuh-Nung Jan, UCSF, USA), rat-anti-Dpn (1:150, Abcam, ab195173), rabbit anti-Dpn (1:800, Yuh-Nung Jan, UCSF, USA), mouse-anti-Pros (1:20, DSHB, MR1A), rabbit anti-phosphohistone 3 (1:500, Invitrogen, PA5-17869), mouse-anti-ATPβ (1:200, Abcam, ab14730), rabbit anti-cyclin E (1:200, Santacruz, sc-33748), rabbit-anti-phosphoAMPK (1:200, Cell Signaling, 2535L), rat-anti-Miranda (1:600, Abcam, 197788), rabbit-anti-cleaved Caspase 3 (1:100, Cell Signaling, D175), rabbit-anti-Cytochrome C (1:200, Cell Signaling, 4272S). The following Alexa Fluor-coupled secondary antibodies (Invitrogen) were used at a 1:1000 dilution: anti-chicken 488, anti-mouse 568/647, anti-rabbit 488/568/647, and anti-rat 488/568/647.

### ROS detection in live third instar larval brain by DHE fluorescence

Third instar larval brains were dissected in Schneider’s media, and each of the brain lobes was carefully cut at the outer margin of the optic lobe to facilitate the penetration of dihydroethidium (DHE) dye (Invitrogen, D1168) into brain tissue.

Subsequently, the brains were incubated in 30 µM DHE for 15 minutes at RT and washed with Schneider’s media for 10 minutes before being mounted in a poly-L-lysine (Sigma, P8920) coated LabTEK chamber (Thermofischer scientific, 155380), immersed in Schneider’s media. The coverslip was placed over the brain very gently to avoid movement. Imaging was performed immediately using the Zeiss LSM710 confocal system with a 63X/1.4 NA Oil objective and excitation/emission of 561nm/605nm.

### TMRM uptake for mitochondrial membrane potential

Larval brains were dissected out in Schneider’s media and treated with 1:150 D(+) Limonene (Lobachemie, 0441600500) (diluted in Schneider’s media) for 4 minutes. Subsequently, brains were immersed in 100 nM Tetramethylrhodamine methyl ester (TMRM) dye (Thermofischer Scientific, T668) for 30 minutes. The brains were washed with Schneider’s media for 5 minutes and mounted in a poly-L-lysine coated LabTEK chamber, immersed in Schneider’s media. Imaging was performed immediately using the Zeiss LSM710 confocal system with a 63X/1.4 NA Oil objective and excitation/emission of 561nm/605nm.

### EdU uptake and detection assay for cell proliferation analysis

EdU assay was performed by using a Click-iTTM EdU Alexa FluorTM 647 imaging kit (Invitrogen, C10340). Third instar larvae were dissected in Schneider’s medium and immediately treated with 10μM EdU (5-ethynyl-2’-deoxyuridine) for 15 minutes at RT. EdU incorporation reaction in DNA was stopped by doing fixation of brains in 4% PFA for 25 minutes. The brains were washed in 0.1% PBST for 30 minutes at room temperature (RT) and blocked with 1% BSA in 0.1% PBST for 1 hour at RT. Following this, the brains were incubated with the required primary antibodies overnight (∼14 hours) at 4D on a shaker. The next day, brains were washed for 20 minutes with 0.1% PBST. Subsequently, the brains were incubated with secondary antibodies against primary antibodies for 1 hour at RT, followed by 20 minutes of washing with 0.1% PBST. Subsequently, the brains were washed for 20 minutes with 0.1% PBST. A Click-iT reaction cocktail was prepared according to the manufacturer’s instructions. The brains were incubated in it for 30 minutes at RT, followed by 10 minutes of washing with 0.1%PBST. For DNA staining, the brains were treated with Hoechst (1:1000, Invitrogen, H3570) in 0.1%PBST for 10 minutes and then washed for 10 minutes with 0.1% PBST before being mounted in Slow-Fade Gold. Subsequent imaging was performed using a Zeiss LSM710 with a 40x/1.4NA oil objective.

### ATP dye uptake assay

3^rd^ instar larval brains were dissected out in Schneider’s media and treated with 1:150 D(+) Limonene (Lobachemie) (diluted in Scheiders media) for 4 minutes. Subsequently, brains were immersed in 30μM ATP-red dye (Merk Millipore, SCT045) for 30 minutes. Afterward, the brains were washed with Schneider’s media for 5 minutes and mounted in a poly-L-lysine coated LabTEK chamber, immersed in Schneider’s media. Imaging was done immediately using the Zeiss LSM710 with 63X/1.4 NA Oil objective using 561 nm laser.

### Microscopy

#### Imaging of fixed samples

Fixed samples were imaged at room temperature using a Zeiss LSM710 or LSM780 inverted confocal microscope at the IISER Pune microscopy facility. A Plan-Apochromat 40x (1.4 NA) or 63x (1.4 NA) oil objective was used for image acquisition. Images were captured at 1024×1024 pixels with an averaging of 2 and an acquisition speed of 8 using Zen2010 software. Fluorescence intensity was maintained within the 8-bit scale (0–255). During confocal microscopy, Hoechst, a DNA dye, was excited using a Diode laser. Argon laser (488 nm) was employed for Alexa Fluor 488, while a DPSS laser was used to image Alexa Fluor 568. Finally, Alexa Alexa Fluor 647 signals were acquired using a 633 nm HeNe laser. Representative images in Fig.s depict type II lineage, chosen to display the maximum number of cells within that lineage.

### Live imaging of type II neuroblast in larval brain

For live imaging of type II neuroblasts, third instar larval brains were dissected at room temperature in Schneider’s *Drosophila* media. An inverted ZMP LSM710 confocal microscope was used to scan the brains on a 35 mm dish in Schneider’s media supplemented with 10% FBS and 10 mM insulin (I0516, Sigma) at 3-minute intervals for three hours at 25°C.

### Transmission electron microscopy of larval neuroblasts

Third instar larval brains were dissected in Schneider’s *Drosophila* media at room temperature. Brains were immediately fixed in 3% glutaraldehyde (Sigma, G5882) for 2hr at 4°C. Washed twice with 0.1 M Sodium cacodylate buffer, pH 7.4, for 10 min each. Then, tissues were further fixed in 1 % Osmium tetroxide (OsO_4_) for 1 hr at 4^0^C. Brains were en-block stained with 2% uranyl acetate and dehydrated for ten minutes each in an increasing alcohol series. Araldite resins A and B were used to infiltrate the tissues, and Araldite resin B was used to embed the tissues and polymerize them for 48 hours at 60^0^C. Using an ultra-microtome (LeicaUC7), ultrathin brain slices of 70 nm were cut and collected on copper slot grids. Lead citrate was used to contrast the sections. The transmission electron microscope, JEM 1400 Plus, JEOL (Japan), at 120 kV, was used to take the images. Neuroblasts were identified due to their distinct large size in the central brain.

### Image quantification and statistical analysis

#### Mitotic index quantification of neuroblast in third instar larval brain

Neuroblasts in the central brain hemisphere were quantified using Miranda as a cortical marker in conjunction with mCD8-GFP<*wor*-Gal4 across various genotypes. The percentage of pH3-positive neuroblasts was then determined and plotted to assess neuroblastproliferation. Type II neuroblastmitotic index was calculated by counting pH3-positive mCD8-GFP<pnt-Gal4 cells within each brain lobe. Statistical significance was assessed using a non-parametric Mann-Whitney t-test

### Analysis of mINPs, GMCs, and neurons number in the type II neuroblast lineage

Dpn-positive cells were classified as mINPs. Ase marked mINPs and GMCs. Dpn-negative, Ase-positive cells were counted as GMCs within each type II neuroblast lineage. Young neurons, characterized by high nuclear Pros expression at the lineage base (59, 60), were quantified from all type II neuroblast lineage Z-sections.

The lineage boundary was delineated by anti-GFP immunostaining to enhance the mCD8-GFP signal. ImageJ’s brightness and contrast tools further ensured that only cells belonging to the same lineage were counted. Cell counts were plotted using GraphPad Prism, and statistical analysis was performed with a non-parametric Mann-Whitney t-test.

### EdU incorporation analysis

Using the ImageJ cell counter, we quantified EdU-positive cells across different planes within single type II lineages for various genotypes. Proliferating mINPs were specifically identified by co-localization of Dpn and EdU. Cell counts were then plotted using GraphPad Prism software, and a non-parametric Mann-Whitney t-test was applied for statistical analysis.

### Nuclear Cyclin E Intensity Measurement

We quantified nuclear Cyclin E intensity in GFP-positive type II neuroblasts. Using ImageJ’s freehand tool, we delineated each neuroblast’s nuclear region of interest. The nuclear to cytoplasmic Cyclin E fluorescence intensity ratio was then calculated for all genotypes. These ratios were plotted using GraphPad Prism software, and a non-parametric Mann-Whitney t-test was applied for statistical analysis.

### Quantification of ATP, ROS and MMP in type II neuroblasts

We measured ATP by ATP-red fluorescence, ROS by quantifying DHE fluorescence and MMP by TMRM fluorescence intensity in type II neuroblasts (mCD8-GFP positive) across different genotypes. For comparison, we also analyzed neighboring wild-type, type I neuroblasts (mCD8-GFP negative) within the same samples. Using ImageJ’s freehand tool, we defined a region of interest (cytoplasmic area) for each neuroblast. The ratio of DHE fluorescence intensity of the GFP-positive neuroblast to a neighboring GFP-negative neuroblast (in the same optical plane) was calculated for all genotypes. These ratios were then plotted using GraphPad Prism software, and a non-parametric Mann-Whitney t-test was performed for statistical analysis.

## Supplementary Figure legends

**Fig. S1. Complex I depletion leads to developmental defects and decrease in type II neuroblast division**

**(A)** The table shows developmental phenotypes of ETC complex I subunits knockdown with different neuronal Gal4s. *wor*-Gal4 and *pnt*-Gal4 drivers were crossed with RNAi lines against subunits of ETC complex I at 29°C and lethality or behavioural phenotype was recorded in the F1 adults. *wor*-Gal4 is expressed in all neuroblasts, and *pnt*-Gal4 is expressed in type II neuroblasts. *wor*-Gal4 driven depletion of complex I showed lethality of animals at different stages of development. But *pnt*-Gal4 driven downregulation of ETC complexes showed a less severe phenotype; animals emerged from pupae, but they were sluggish and died within 10-15 days after eclosion. (**B)** Representative images of snapshots during live imaging of type II neuroblasts of genotypes, mCherry^i^ (n=15 neuroblast, 3 brains) and ND-51^i^(25,5). Control, mCherry^i^ type II neuroblasts divided at least once during a live imaging period of 40 minutes, but ND-51^i^ neuroblasts failed to do so. Yellow dashed circle marks type II neuroblast. Before starting of division, four cells of the lineage labelled as 1,2,3 and 4. After the completion of one division, a newly generated cell labelled as 5 is seen.

**Fig. S2. Analysis of caspase and differentiation in different RNAi lines and Gal4 dilution effects in complex I depleted type II neuroblast lineages**

**(A)** Representative images of cleaved caspase 3 immunostaining (red) in type II neuroblast lineage of the indicated genotype. Caspase 3 expression in ND-51^i^ was comparable to mCherry^i^ control. **(B)** Representative images of type II neuroblast lineages of the mentioned genotypes. Dpn positive mINPs (grey). Dpn negative/Ase (red) positive cells GMCs (yellow arrowheads). Scale bar-10µm **(C)** Quantification of mINPs in type II neuroblast lineages of genotypes, mCherry^i^ (n=14 type II neuroblast lineages, 3 brains), ND-51^i^ BL 29534 (26, 6), and mito-GFP;ND-51^i^ (11, 3). **(D)** Quantification of GMCs in type II neuroblast lineages of genotypes, mCherry^i^ (n=15 type II neuroblast lineages, 3 brains), ND-51^i^ BL 29534 (20, 6), and mito-GFP;ND-51^i^ (11, 3). **(E)** Representative images of neurons stained with high nuclear Pros (red) in type II neuroblast lineages of indicated genotypes. Scale bar-10µm. **(F)** Quantification of neurons in type II neuroblast lineages of genotypes, mCherry^i^ (n=12 type II neuroblast lineages, 3 brains), ND-51^i^ BL 29534 (17, 6), and mito-GFP;ND-51^i^ (10, 3).

**Fig. S3. mINPs were decreased in mitosis in complex I depleted type II neuroblast lineages**

**(A)** Representative images for pH3 labelled (grey) mINPs in the type II neuroblast lineage of the mentioned genotypes. Dpn positive mINPs (red). **(B)** Quantification of pH3 positive mINPs in type II neuroblast lineages of genotypes, mCherry^i^ (n=13 type II neuroblast lineages, 5 brains), Drp1^SD^;mCherry^i^ (20, 5), ND-51^i^ (5, 4), Drp1^SD^;ND-51^i^ (12, 3).

**Fig. S4. ATP levels and pAMPK levels were comparable to controls in complex I depleted type II neuroblast lineages**

**(A)** Representative images of ATP levels in type II neuroblast using ATP-Red dye (red). mCD8-GFP marks the boundary of neuroblasts and lineage cells (green). White dashed circle label type II neuroblast and surrounding non-mutant type I neuroblast shown by yellow arrowheads. Scale bar-10μm. **(B)** Quantification of ATP-Red dye intensity normalised to surrounding type I neuroblast in genotypes, mCherry^i^ (22 neuroblast, 3 brain) and ND-51^i^ (24,3). **(C)** Representative figure of pAMPK immunostaining (red) for ATP detection in type II neuroblast. mCD8-GFP marks the boundary of neuroblasts and lineage cells (green). White dashed circle label type II neuroblast. Scale bar-10μm. **(D)** Quantification of phospho-AMPK intensity in type II neuroblast of genotypes, mCherry^i^ (10 neuroblast, 3 brain) and ND-51^i^ (6,3).

**Fig. S5. Analysis of ROS, membrane potential and mitochondrial morphology in complex I depleted type II neuroblast lineages**

**(A)** Representative images of DHE fluorescence in type II neuroblasts of the mentioned genotype. White dashed circle denotes type II neuroblast. mCD8-GFP marks the lineage cell (green). Surrounding type I neuroblasts are shown by yellow arrowheads. **(B)** Quantification of DHE intensity in type II neuroblasts normalized to surrounding type I neuroblasts of the genotypes, mCherry^i^ (9 type II neuroblasts, 3 brains), SOD1^i^ (16, 3), and SOD2^i^ (11, 3). **(C)** Representative images showing DHE fluorescence in type II neuroblasts of the mentioned genotype. **(D)** Quantification of DHE fluorescence in type II neuroblast normalized to surrounding type I neuroblast of the following genotypes: mCherry^i^ (9 type II neuroblasts, 3 brains), SOD2^OE^ (14,4), ND-51^i^ (30, 5), and SOD2^OE^;ND-51^i^ (31,4). **(E)** Representative images depicting TMRM fluorescence (red, circled by white dashed line) in type II neuroblasts. Surrounding control type I neuroblasts are shown by yellow arrowheads. Scale bar-10µm. **(F)** Quantification of TMRM fluorescence intensity in type II neuroblast, as compared to the intensity of neighbouring type I neuroblast. The following genotypes were assessed: mCherry^i^ (n=18 type II neuroblasts, 5 brains), SOD1^i^ (19,5), SOD2^i^ (27,7), and SOD2^OE^;ND-51^i^ (10,4). **(G)** Representative images of type II neuroblasts (white dashed outline) and muscle mitochondria stained by using anti-ATPB antibody (green). Knockdown of SOD1 and SOD2 in *Drosophila* type II neuroblasts and larval muscle by using *pnt*-Gal4 and *mhc*-Gal4 respectively, induces mitochondrial fragmentation. Overexpression of SOD2 in ND-51^i^ neuroblasts and muscle restored the fused mitochondrial state compared to ND-51^i^ alone (Fig. 3).

**Fig. S6. G1/S transition defect is alleviated in SOD^OE^ in complex I depleted type II neuroblast lineages**

**(A)** Representative images showing immunostaining of mINPs (Dpn, red) and EdU-positive cells (grey) in the lineages of type II neuroblasts of the mentioned genotypes. mINPs present in S-phase are labelled for both Dpn and EdU. Scale bar-10µm. **(B)** Quantification of total EdU-positive cells (red) in the lineages of type II neuroblasts of genotypes:mCherry^i^ (n=12 type II neuroblast lineages, 4 brains), SOD2^OE^ (12,4), ND-51^i^ (15, 5), and SOD2^OE^;ND-51^i^ (12,5). **(C)** Quantification of percent EdU positive mINPs (Yellow arrowheads) in the lineages of type II neuroblasts of genotypes: mCherry^i^ (n=15 type II neuroblast lineages, 4 brains), SOD2^OE^ (13,4), ND-51^i^ (12, 5), and SOD2^OE^;ND-51^i^ (15,5). **(D)** Representative images of cyclin E immunostaining (red) in type II neuroblasts (encircled by a dashed line) of indicated genotypes. Scale bar-10µm. **(E)** Quantification of cyclin E intensity in nucleus with respect to cytoplasm of type II neuroblasts of following genotypes:: mCherry^i^ (n=17 type II neuroblasts, 3 brains) and SOD2^OE^;ND-51^i^ (20,4).

**Fig. S7. Mitochondrial fusion by Drp1 depletion rescues the differentiation defect in SOD2 depleted type II neuroblasts**

**(A)** Representative images showing immunostaining for mINPs and GMCs of the mentioned genotypes. Dpn positive mINPs (grey). Ase positive (red)/Dpn negative cells GMCs (yellow arrowheads). **(B)** Quantification of mINPs in type II neuroblast lineages of genotype: mCherry^i^ (n=25 type II neuroblast lineages, 5 brains), and SOD2^i^ (17, 4), and Drp1^SG^;SOD2^i^ (17,4). **(C)** Quantification of GMCs in type II neuroblast lineages of genotypes: mCherry^i^ (n=24 type II neuroblast lineages, 5 brains), SOD2^i^ (17, 4), and Drp1^SG^;SOD2^i^ (15,4). **(D)** Representative images of immunostaining of neurons with high nuclear Pros (red) in type II neuroblast lineages of indicated genotypes. Scale bar-10µm. **(E)** Quantification of neurons in type II neuroblast lineages of genotypes: mCherry^i^ (n=19 type II neuroblast lineages, 5 brains), SOD2^i^ (14, 4), and Drp1^SG^;SOD2^i^ (11,3). Data for mCherry^i^ and SOD2^i^ are repeated from Fig. 6.

## REFERENCES

1. M. J. Son, B. R. Jeong, Y. Kwon, Y. S. Cho, Interference with the mitochondrial bioenergetics fuels reprogramming to pluripotency via facilitation of the glycolytic transition. Int. J. Biochem. Cell Biol. 45, 2512–2518 (2013).

2. W. Liu, et al., Mitochondrial metabolism transition cooperates with nuclear reprogramming during induced pluripotent stem cell generation. Biochem. Biophys. Res. Commun. 431, 767–771 (2013).

3. S. M. Schieke, et al., Mitochondrial metabolism modulates differentiation and teratoma formation capacity in mouse embryonic stem cells. J. Biol. Chem. 283, 28506–28512 (2008).

4. S. Varum, et al., Enhancement of human embryonic stem cell pluripotency through inhibition of the mitochondrial respiratory chain. Stem Cell Res. 3, 142– 156 (2009).

5. C. C. F. Homem, M. Repic, J. A. Knoblich, Proliferation control in neural stem and progenitor cells. Nat. Rev. Neurosci. 16, 647–659 (2015).

6. D. Dubal, P. Moghe, R. K. Verma, B. Uttekar, R. Rikhy, Mitochondrial fusion regulates proliferation and differentiation in the type II neuroblast lineage in Drosophila. PLoS Genet. 18, e1010055 (2022).

7. J. van den Ameele, A. H. Brand, Neural stem cell temporal patterning and brain tumour growth rely on oxidative phosphorylation. Elife 8 (2019).

8. F. Bonnay, et al., Oxidative metabolism drives immortalization of neural stem cells during tumorigenesis. Cell 182, 1490–1507.e19 (2020).

9. S. Zhu, S. Barshow, J. Wildonger, L. Y. Jan, Y.-N. Jan, Ets transcription factor Pointed promotes the generation of intermediate neural progenitors in Drosophila larval brains. Proc. Natl. Acad. Sci. U. S. A. 108, 20615–20620 (2011).

10. B. C. Bello, N. Izergina, E. Caussinus, H. Reichert, Amplification of neural stem cell proliferation by intermediate progenitor cells in Drosophila brain development. Neural Dev. 3, 5 (2008).

11. J. A. Knoblich, Asymmetric cell division: recent developments and their implications for tumour biology. Nat. Rev. Mol. Cell Biol. 11, 849–860 (2010).

12. C. Kelsom, W. Lu, Uncovering the link between malfunctions in Drosophila neuroblast asymmetric cell division and tumorigenesis. Cell Biosci. 2, 38 (2012).

13. C. J. Garcia, J. Khajeh, E. Coulanges, E. I.-J. Chen, E. Owusu-Ansah, Regulation of mitochondrial complex I biogenesis in Drosophila flight muscles. Cell Rep. 20, 264–278 (2017).

14. A.-N. A. Agip, I. Chung, A. Sanchez-Martinez, A. J. Whitworth, J. Hirst, Cryo-EM structures of mitochondrial respiratory complex I from Drosophila melanogaster. Elife 12, e84424 (2023).

15. V. Guarani, et al., TIMMDC1/C3orf1 functions as a membrane-embedded mitochondrial complex I assembly factor through association with the MCIA complex. Mol. Cell. Biol. 34, 847–861 (2014).

16. I. Berger, et al., Mitochondrial complex I deficiency caused by a deleterious NDUFA11 mutation. Ann. Neurol. 63, 405–408 (2008).

17. D. A. Stroud, et al., Accessory subunits are integral for assembly and function of human mitochondrial complex I. Nature 538, 123–126 (2016).

18. H. Deng, S. Takashima, M. Paul, M. Guo, V. Hartenstein, Mitochondrial dynamics regulates Drosophila intestinal stem cell differentiation. Cell Death Discov. 4, 17 (2018).

19. S. Chowdhary, D. Tomer, D. Dubal, D. Sambre, R. Rikhy, Analysis of mitochondrial organization and function in the Drosophila blastoderm embryo. Sci. Rep. 7, 5502 (2017).

20. T. Trevisan, et al., Manipulation of mitochondria dynamics reveals separate roles for form and function in mitochondria distribution. Cell Rep. 23, 1742–1753 (2018).

21. S. Mandal, W. A. Freije, P. Guptan, U. Banerjee, Metabolic control of G1-S transition: cyclin E degradation by p53-induced activation of the ubiquitin-proteasome system. J. Cell Biol. 188, 473–479 (2010).

22. E. Owusu-Ansah, A. Yavari, S. Mandal, U. Banerjee, Distinct mitochondrial retrograde signals control the G1-S cell cycle checkpoint. Nat. Genet. 40, 356– 361 (2008).

23. D. J. Parker, et al., A new mitochondrial pool of cyclin E, regulated by Drp1, is linked to cell-density-dependent cell proliferation. J. Cell Sci. 128, 4171–4182 (2015).

24. K. Mitra, R. Rikhy, M. Lilly, J. Lippincott-Schwartz, DRP1-dependent mitochondrial fission initiates follicle cell differentiation during Drosophila oogenesis. J. Cell Biol. 197, 487–497 (2012).

25. C.-H. Yao, et al., Mitochondrial fusion supports increased oxidative phosphorylation during cell proliferation. Elife 8 (2019).

26. N. Ikon, R. O. Ryan, Cardiolipin and mitochondrial cristae organization. Biochim. Biophys. Acta Biomembr. 1859, 1156–1163 (2017).

27. R. W. Gilkerson, J. M. L. Selker, R. A. Capaldi, The cristal membrane of mitochondria is the principal site of oxidative phosphorylation. FEBS Lett. 546, 355–358 (2003).

28. F. Vogel, C. Bornhövd, W. Neupert, A. S. Reichert, Dynamic subcompartmentalization of the mitochondrial inner membrane. J. Cell Biol. 175, 237–247 (2006).

29. D. H. Song, et al., Biophysical significance of the inner mitochondrial membrane structure on the electrochemical potential of mitochondria. Phys. Rev. E Stat. Nonlin. Soft Matter Phys. 88, 062723 (2013).

30. R. Willows, et al., Phosphorylation of AMPK by upstream kinases is required for activity in mammalian cells. Biochem. J. 474, 3059–3073 (2017).

31. L. K. Sharma, et al., Mitochondrial respiratory complex I dysfunction promotes tumorigenesis through ROS alteration and AKT activation. Hum. Mol. Genet. 20, 4605–4616 (2011).

32. J. Ježek, K. Cooper, R. Strich, Reactive oxygen species and mitochondrial dynamics: The yin and yang of mitochondrial dysfunction and cancer progression. Antioxidants (Basel) 7, 13 (2018).

33. M. Khacho, et al., Mitochondrial dynamics impacts stem cell identity and fate decisions by regulating a nuclear transcriptional program. Cell Stem Cell 19, 232–247 (2016).

34. E. Owusu-Ansah, U. Banerjee, Reactive oxygen species prime Drosophila haematopoietic progenitors for differentiation. Nature 461, 537–541 (2009).

35. M. P. Murphy, How mitochondria produce reactive oxygen species. Biochem. J. 417, 1–13 (2009).

36. C. A. Juan, J. M. Pérez de la Lastra, F. J. Plou, E. Pérez-Lebeña, The chemistry of reactive oxygen species (ROS) revisited: Outlining their role in biological macromolecules (DNA, lipids and proteins) and induced pathologies. Int. J. Mol. Sci. 22, 4642 (2021).

37. D. Han, F. Antunes, R. Canali, D. Rettori, E. Cadenas, Voltage-dependent anion channels control the release of the superoxide anion from mitochondria to cytosol. J. Biol. Chem. 278, 5557–5563 (2003).

38. E. Cadenas, K. J. Davies, Mitochondrial free radical generation, oxidative stress, and aging. Free Radic. Biol. Med. 29, 222–230 (2000).

39. M. D. Brand, The sites and topology of mitochondrial superoxide production. Exp. Gerontol. 45, 466–472 (2010).

40. L. K. Sharma, J. Lu, Y. Bai, Mitochondrial respiratory complex I: structure, function and implication in human diseases. Curr. Med. Chem. 16, 1266–1277 (2009).

41. V. Ten, A. Galkin, Mechanism of mitochondrial complex I damage in brain ischemia/reperfusion injury. A hypothesis. Mol. Cell. Neurosci. 100, 103408 (2019).

42. A. J. Lambert, M. D. Brand, Superoxide production by NADH:ubiquinone oxidoreductase (complex I) depends on the pH gradient across the mitochondrial inner membrane. Biochem. J. 382, 511–517 (2004).

43. M. P. Murphy, Understanding and preventing mitochondrial oxidative damage. Biochem. Soc. Trans. 44, 1219–1226 (2016).

44. A. Kohler, A. Barrientos, F. Fontanesi, M. Ott, The functional significance of mitochondrial respiratory chain supercomplexes. EMBO Rep. 24, e57092 (2023).

45. J. Berndtsson, et al., Respiratory supercomplexes enhance electron transport by decreasing cytochrome c diffusion distance. EMBO Rep. 21, e51015 (2020).

46. G. S. Shadel, T. L. Horvath, Mitochondrial ROS signaling in organismal homeostasis. Cell 163, 560–569 (2015).

47. R. A. Haworth, D. R. Hunter, The Ca2+-induced membrane transition in mitochondria. II. Nature of the Ca2+ trigger site. Arch. Biochem. Biophys. 195, 460–467 (1979).

48. K. D. Garlid, A. D. Beavis, Evidence for the existence of an inner membrane anion channel in mitochondria. Biochim. Biophys. Acta 853, 187–204 (1986).

49. D. B. Zorov, C. R. Filburn, L. O. Klotz, J. L. Zweier, S. J. Sollott, Reactive oxygen species (ROS)-induced ROS release: a new phenomenon accompanying induction of the mitochondrial permeability transition in cardiac myocytes. J. Exp. Med. 192, 1001–1014 (2000).

50. M. J. Barsoum, et al., Nitric oxide-induced mitochondrial fission is regulated by dynamin-related GTPases in neurons. EMBO J. 25, 3900–3911 (2006).

51. X. Chang, et al., ROS-Drp1-mediated mitochondria fission contributes to hippocampal HT22 cell apoptosis induced by silver nanoparticles. Redox Biol. 63, 102739 (2023).

52. V. Brillo, L. Chieregato, L. Leanza, S. Muccioli, R. Costa, Mitochondrial dynamics, ROS, and cell signaling: A blended overview. Life (Basel) 11, 332 (2021).

53. Q. Huang, et al., Increased mitochondrial fission promotes autophagy and hepatocellular carcinoma cell survival through the ROS-modulated coordinated regulation of the NFKB and TP53 pathways. Autophagy 12, 999–1014 (2016).

54. M. C. Ludikhuize, et al., Mitochondria define intestinal stem cell differentiation downstream of a FOXO/Notch axis. Cell Metab. 32, 889–900.e7 (2020).

55. S. Chowdhary, S. Madan, D. Tomer, M. Mavrakis, R. Rikhy, Mitochondrial morphology and activity regulate furrow ingression and contractile ring dynamics in Drosophila cellularization. Mol. Biol. Cell 31, 2331–2347 (2020).

56. A. Öztürk-Çolak, et al., FlyBase: updates to the Drosophila genes and genomes database. Genetics 227 (2024).

57. V. K. Jenkins, A. Larkin, J. Thurmond, FlyBase Consortium, Using FlyBase: A database of Drosophila genes and genetics. Methods Mol. Biol. 2540, 1–34 (2022).

58. J. Zirin, et al., Large-scale transgenic Drosophila resource collections for loss-and gain-of-function studies. Genetics 214, 755–767 (2020).

59. O. A. Bayraktar, J. Q. Boone, M. L. Drummond, C. Q. Doe, Drosophila type II neuroblast lineages keep Prospero levels low to generate large clones that contribute to the adult brain central complex. Neural Dev. 5, 26 (2010).

60. T. J. Samuels, et al., Neuronal upregulation of Prospero protein is driven by alternative mRNA polyadenylation and Syncrip-mediated mRNA stabilisation. Biol. Open 9, bio049684 (2020).

