## Supplementary figures and images for "Electron transport chain complex I and mitochondrial fusion regulate ROS for differentiation in *Drosophila* neural stem cells"

### Figure S1

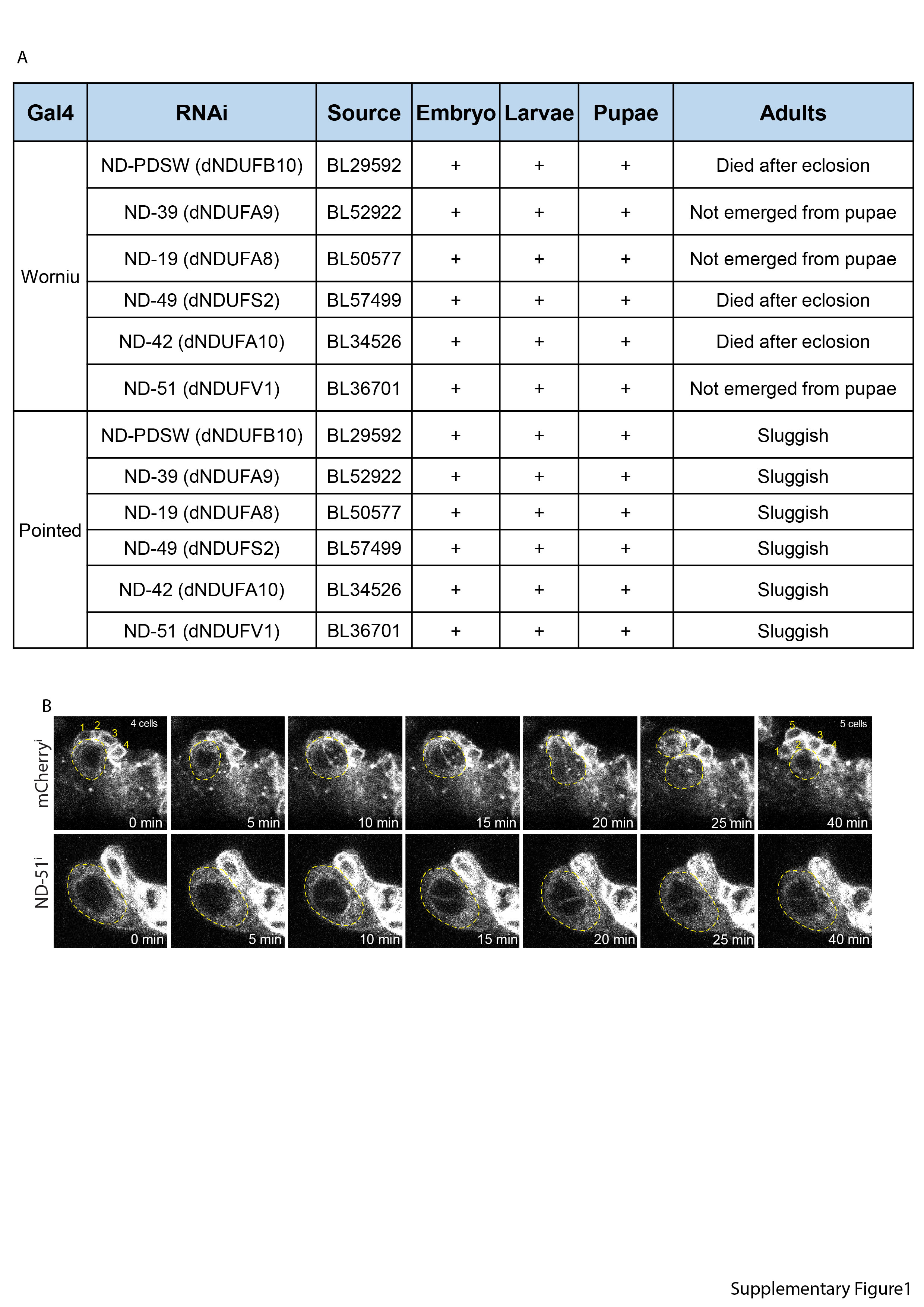

### Figure S2

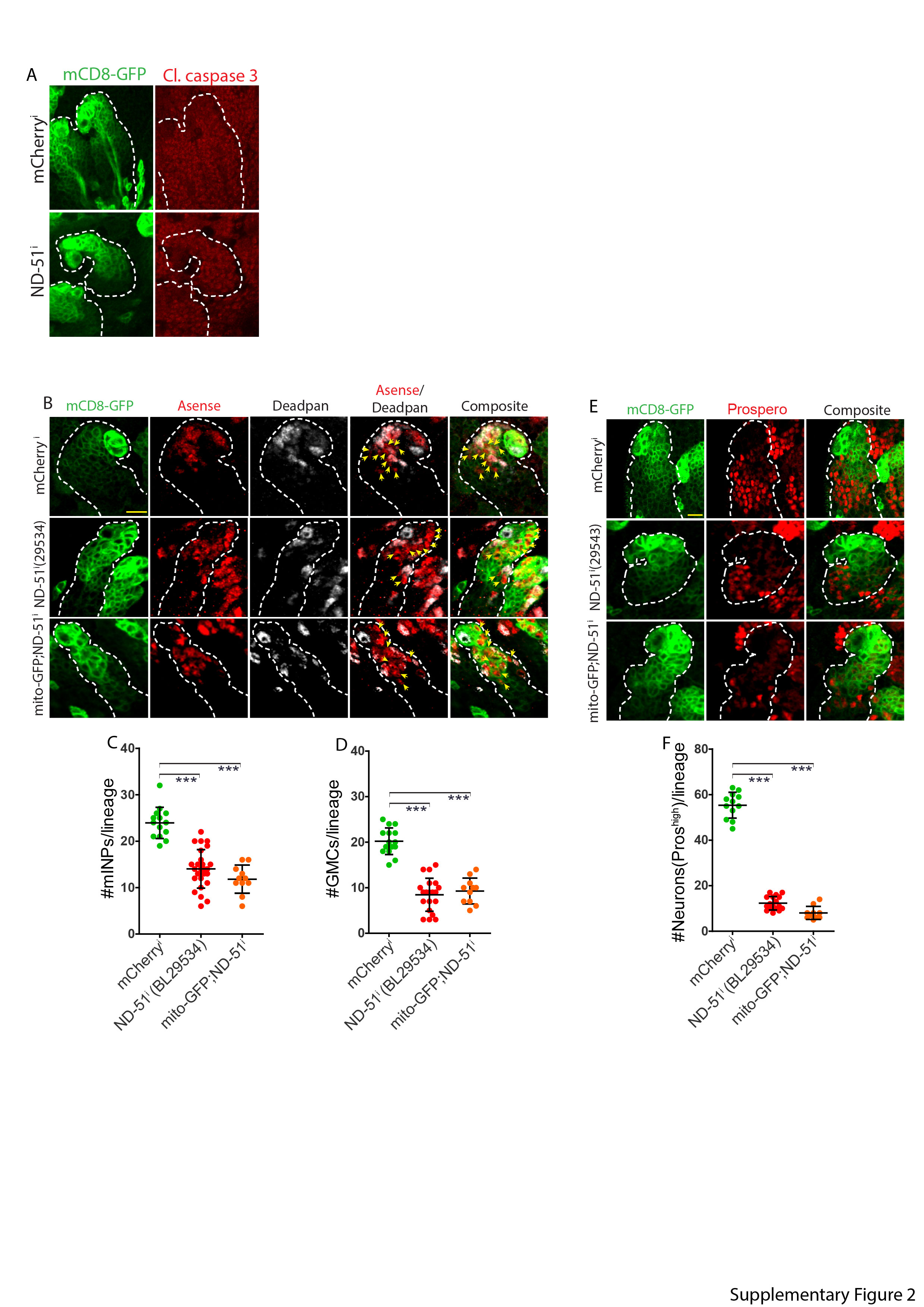

### Figure S3

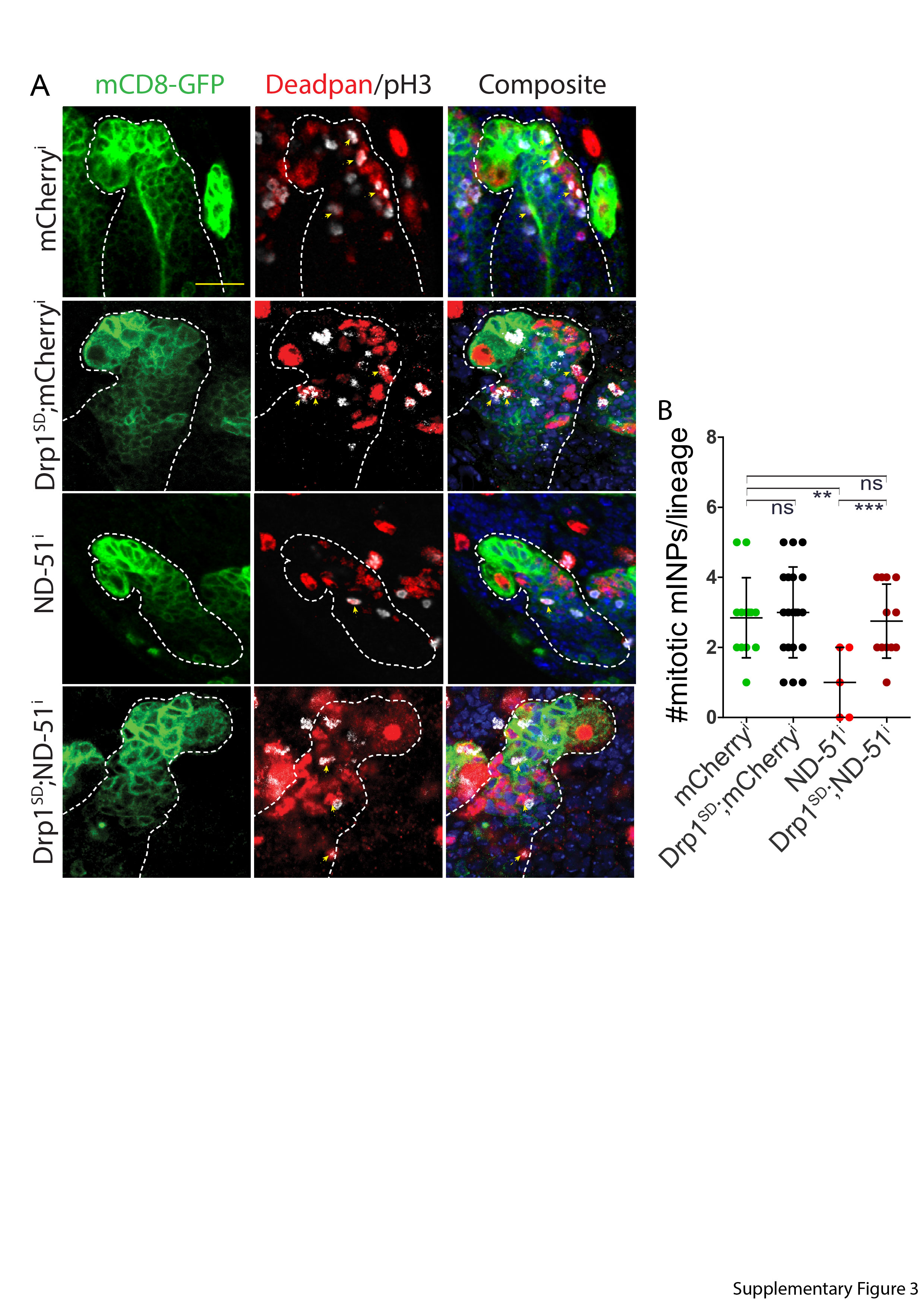

### Figure S4

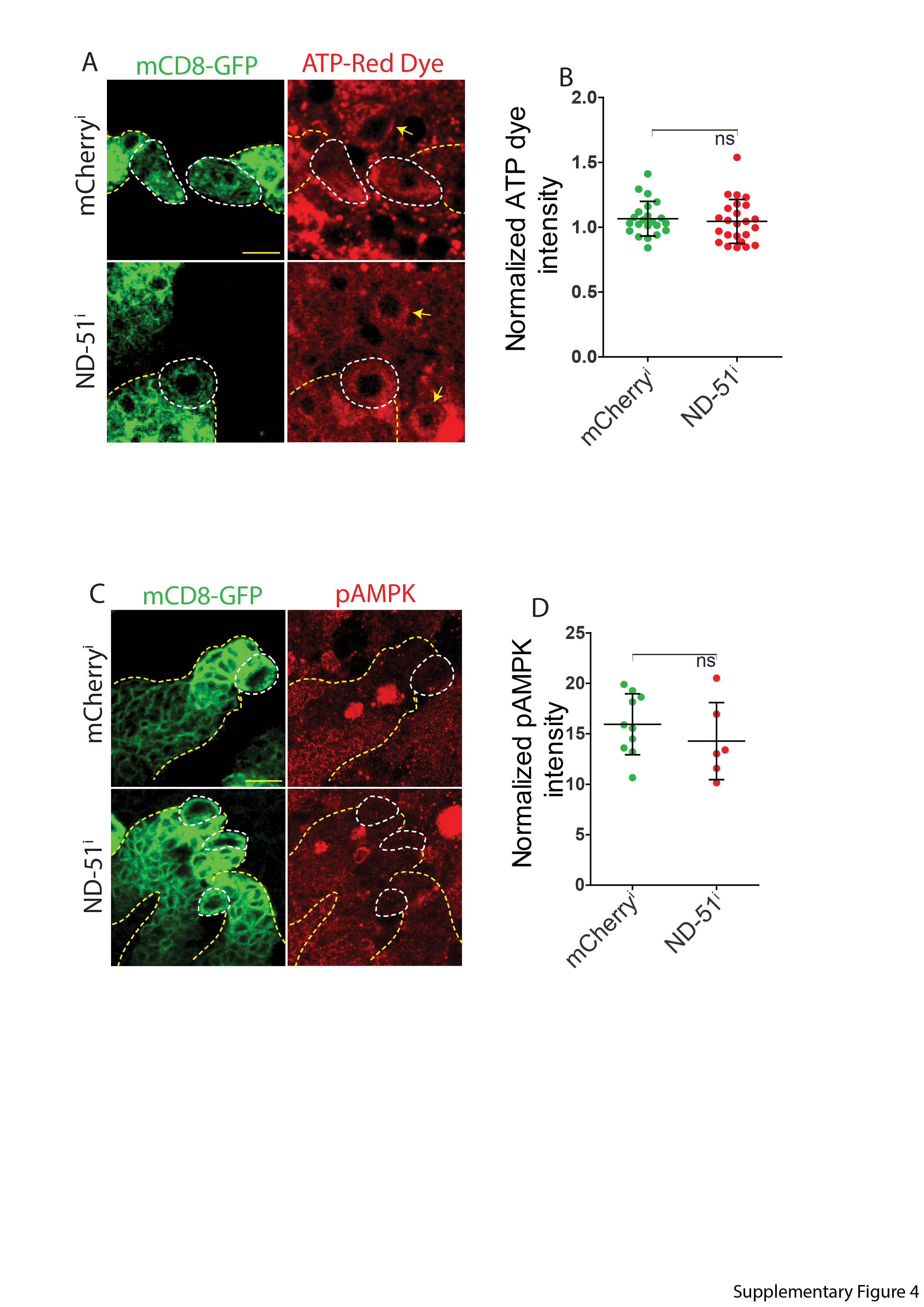

### Figure S5

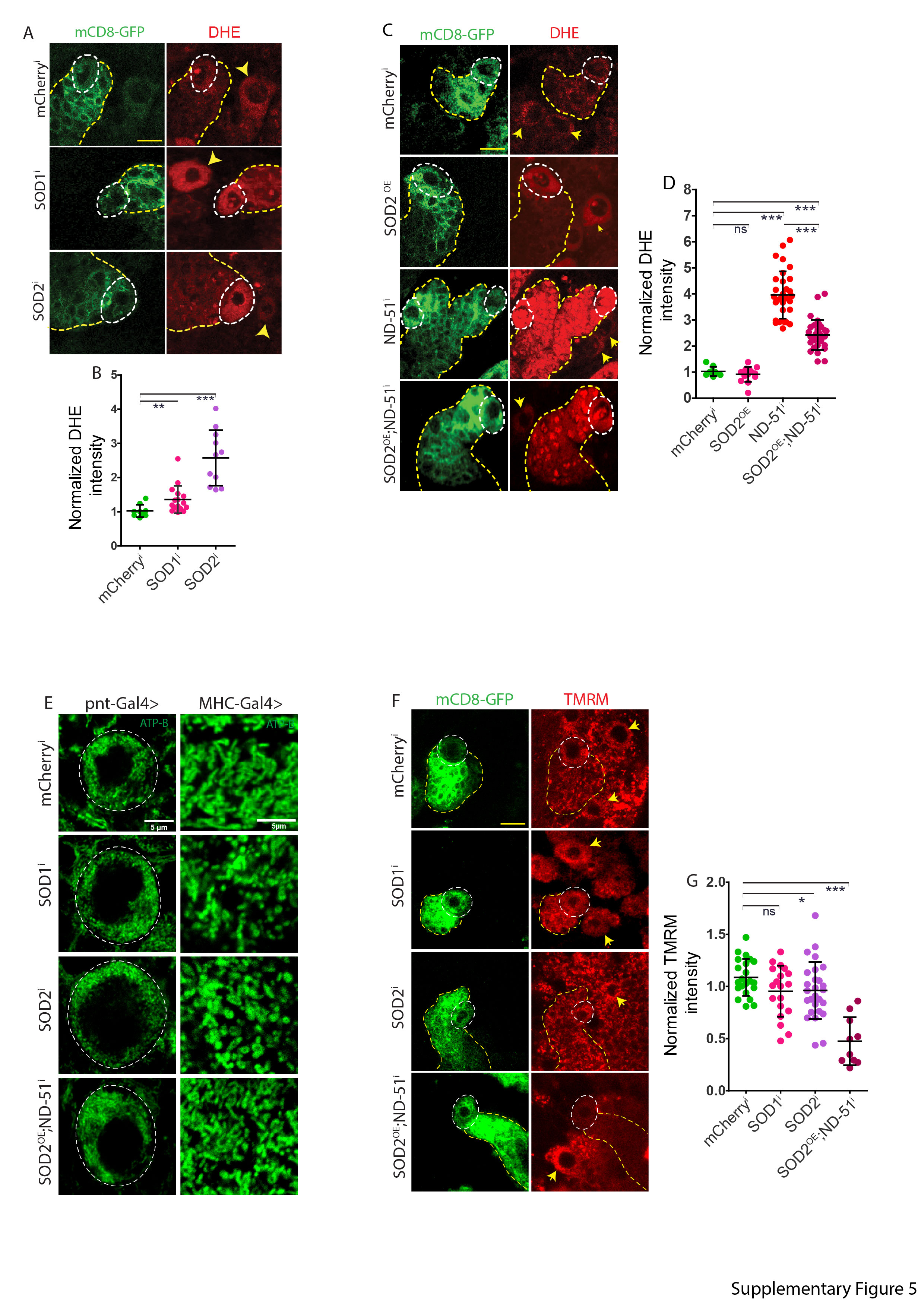

### Figure S6

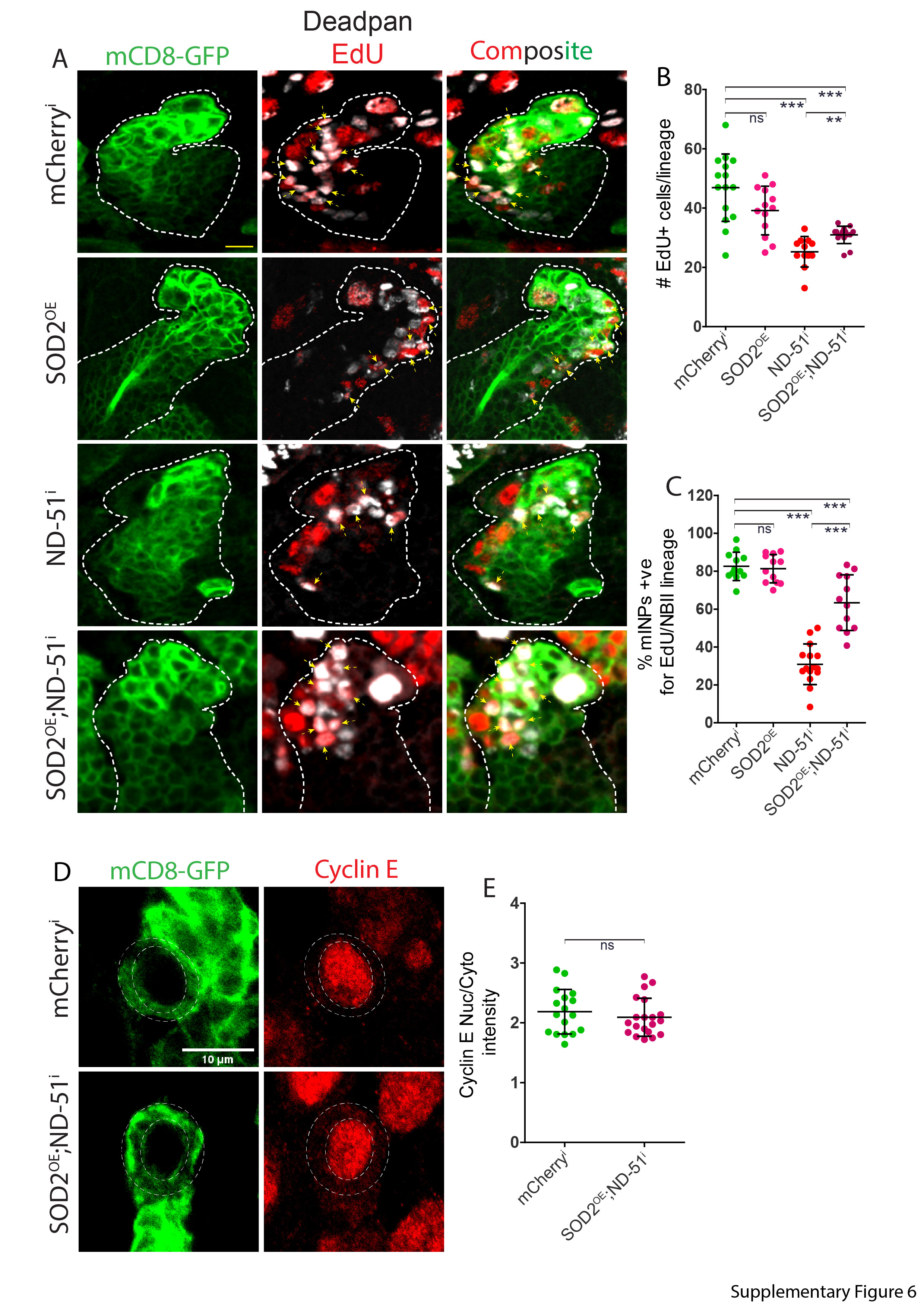

### Figure S7

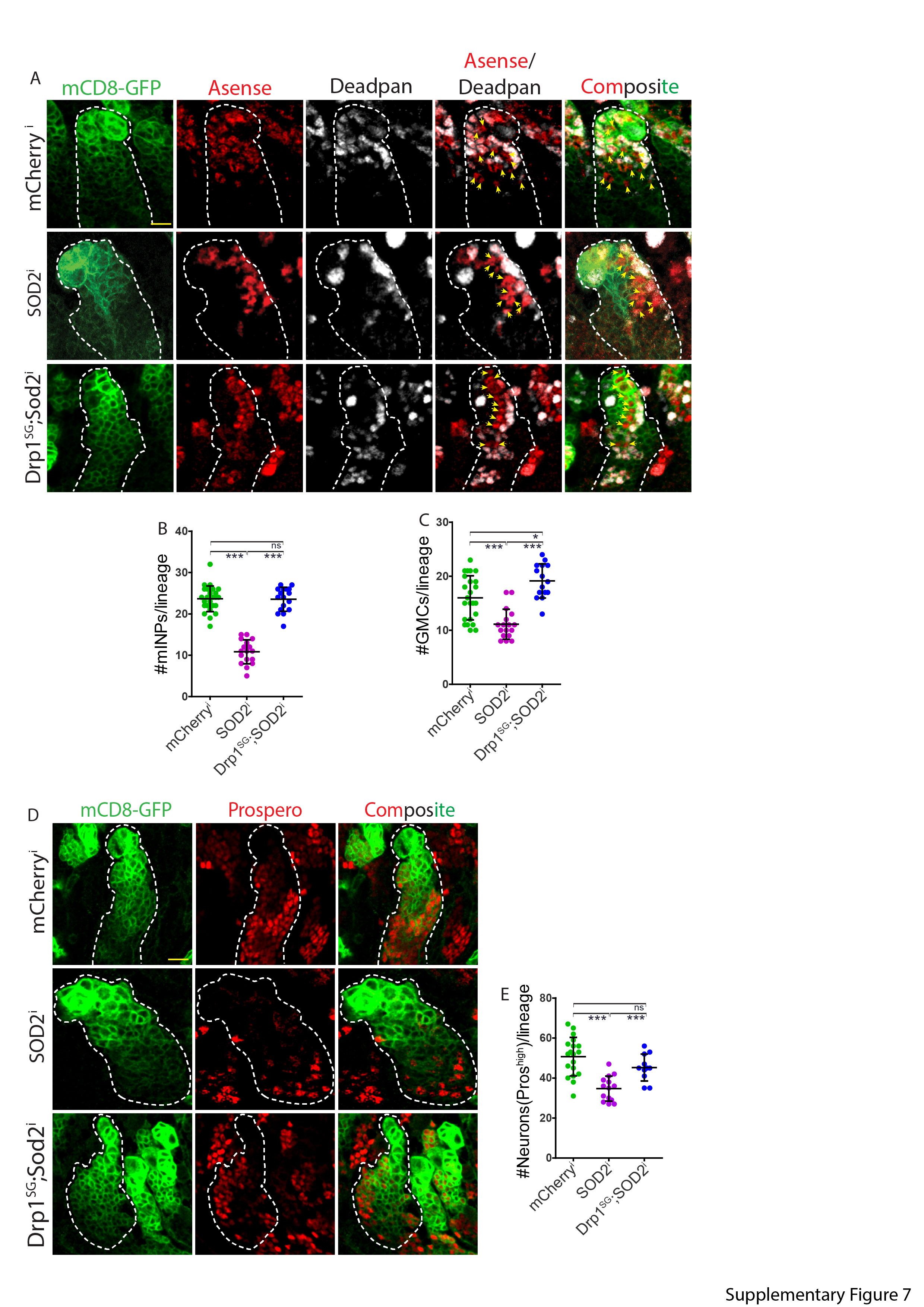
